# Shared roles and team membership are reflected in functional connectome similarity: Neural evidence from real-world volleyball teams

**DOI:** 10.64898/2026.05.09.723964

**Authors:** Jer-Jen Chang, Yu-Chun Chen, Yen-Sheng Chiang

## Abstract

In task-oriented teams, long-term coordination among specialized roles may contribute to shared patterns of cognition and behavior, yet little is known about how such experience is reflected in brain functional organization. Here, we examined whether cross-individual differences in whole-brain functional connectivity were associated with court position and team membership in professional volleyball players. In the resting-state and naturalistic volleyball game-viewing conditions, we analyzed dyadic functional connectivity differences to test whether effects of shared position and team were evident across intrinsic and contextually engaged brain states, controlling for differences in playing time and performance-related statistics. We found that same-position players showed smaller functional connectivity differences. These effects were most prominent and widespread across brain networks during game viewing, whereas at rest they were specific to the somatomotor network. Team membership was also associated with smaller functional connectivity differences during game viewing, although position × team interactions varied across networks after covariate adjustment. A complementary machine learning classifier further showed that shared position could be predicted from intersubject differences in functional connectivity with accuracy exceeding a frequency-based baseline. Together, these findings suggest that shared role-specific and team-based experience may contribute to structured similarity in functional brain organization within a real-world team setting.

## Introduction

In task-oriented groups, performance is gradually built through long-term coordination among specialized roles. Through repeated interaction, idiosyncratic individuals may develop a sense of shared reality—the experience of holding common internal states (e.g., thoughts, feelings, or beliefs) about the task environment (Rossignac-Milon et al., 2021). In sports psychology, researchers have argued that “team chemistry” may be supported by both overlapping knowledge structures and complementary individual differences among teammates, which together facilitate coordination, cohesion, and effective performance (Eldadi & Tenenbaum, 2025; Filho et al., 2022; Gershgoren et al., 2016). Although these ideas are widely discussed in sports commentary and organizational research, much of the existing evidence is derived from interviews and questionnaire-based measures (Eldadi & Tenenbaum, 2025; Gershgoren et al., 2016; McLaren & Spink, 2016; Tamminen et al., 2016). By comparison, little is known about how individual distinctiveness as well as similarity arising from team-based experience are reflected in the brain, particularly in the context of team sports.

Whole-brain functional connectivity analysis has been widely used to characterize stable, individual-specific patterns of brain functional organization (Beaty et al., 2018; Finn et al., 2015; Mantwill et al., 2022; Miranda-Dominguez et al., 2014). Such cross-individual distinctiveness has been shown to be robust and reliable across both resting-state conditions and naturalistic paradigms, such as movie viewing (Kroll et al., 2023; Vanderwal et al., 2017), earning the characterization of a neural “fingerprint” (Finn et al., 2015). Subsequent research has further demonstrated that individual differences in functional connectivity patterns are state-dependent, varying as a function of the tasks or contexts individuals engage in (Geerligs et al., 2015; Gratton et al., 2018). In addition to task state, longer-term social contexts and cultural experience may also contribute to variability in functional connectivity (Luo et al., 2022). Consistent with this view, a series of studies has shown that people who are closer in social networks exhibit higher neural similarity—that is, smaller differences—both during resting-state scans (Hyon, Youm, et al., 2020) and during naturalistic viewing tasks (Hyon, Kleinbaum, et al., 2020; Parkinson et al., 2018). In other words, although individuals exhibit distinct functional connectivity profiles, the degree of variability among them may not be uniform across social groups. Whether shared experience within highly coordinated real-world teams contributes to such variability remains unclear. Here, we ask whether participation in highly coordinated real-world groups contributes to structured variability in functional connectivity.

We conceptualize volleyball teams as socially organized systems in which individuals occupy differentiated yet interdependent functional roles. Within teams, court position provides a meaningful index of role-specific experience. These role-specific demands may provide a more specific source of similarity than team membership alone. The possibility that role-specific experience is associated with functional brain organization is motivated by findings that long-term sensorimotor experience is associated with altered resting-state functional connectivity. Cross-sectional studies have shown that individuals with long-term athletic experience often exhibit different patterns of resting-state functional connectivity compared with non-athletes (Raichlen et al., 2016; Yang et al., 2020; Zhang et al., 2024; Zhou et al., 2024). Consistent with this, a recent meta-analysis synthesizing data across different sports identified consistent alterations in functional connectivity, particularly within default mode, ventral attention, visual, sensorimotor, and related networks, relative to healthy controls (Yan et al., 2025). Notably, comparisons across different types of elite athletes further suggest sport-specific structural and connectivity adaptations (Sahin Ozarslan & Duru, 2025), and resting-state functional connectivity in motor, default mode, frontoparietal, and attentional networks has also been associated with soccer players’ physical and skill performance (Abbasi et al., 2025). Taken together, these findings suggest that long-term sensorimotor training experience may gradually shape intrinsic brain architecture.

Compared with many interceptive sports in which individual reaction is primary (Fajen et al., 2009; Tresilian, 2004), volleyball requires continuous integration of visual cues, motor planning, and team-based strategy in a dynamic multi-agent environment. Moreover, players in the same position within the same team undergo shared training, coordination, and performance experiences over extended periods. These characteristics make volleyball a compelling naturalistic setting for examining whether role- and team-based experience are reflected in cross-individual similarity in whole-brain functional connectivity.

Using professional volleyball teams as a naturalistic model system, we examine whether cross-individual differences in whole-brain functional connectivity were associated with players’ court position and team membership. We hypothesized that cross-individual differences in functional connectivity patterns would be smaller between players sharing the same position than in different positions, and this effect would be stronger when the players belonged to the same team. We examined cross-individual differences in network-level functional connectivity in both resting-state and naturalistic volleyball game-viewing conditions, allowing us to evaluate whether position- and team-related effects were expressed across intrinsic and contextually engaged brain states. We also additionally controlled for players’ playing time- and performance-related variables. As a complementary analysis, we trained a deep learning model using whole-brain high-dimensional connectivity patterns to predict whether two players occupied the same position.

## Methods

### Participants

Twenty-five male volleyball players from Taiwan’s University Volleyball League (Division I) participated in this study (mean age = 20.6 years, SD = 1.5, range = 18.3–24.2 years). The sample comprised two professional volleyball teams: Team A from the National Taipei University of Education (*n* = 15) and Team B from the National Taiwan University of Sport (*n* = 10). All participants had at least nine years of formal volleyball training experience (mean = 10.36 years, SD = 1.32) and trained an average of 23 hours per week during the competitive season. None reported neurological or psychiatric disorders and all met MRI safety criteria (**Table 1**).

**Table 1.** Demographic and training characteristics of the participants.

|  | All participants<br>(n = 25) | Team A<br>(n = 15) | Team B<br>(n = 10) |
| --- | --- | --- | --- |
| mean age ± SD (years) | 20.6 ± 1.5 | 20.4 ± 1.5 | 21.0 ± 1.4 |
| Years of education | 14.4 ± 1.6 | 14.33 ± 1.7 | 14.50 ± 1.5 |
| Years of volleyball training experience | 10.36 ± 1.3 | 10.47 ± 1.3 | 10.2 ± 1.4 |
| Weekly training hours | 23 | 24 | 22 |
| Right-handed participants (%) | 84.0 | 73.3 | 100 |

Performance data were obtained from official match records within five months of the 2022 season, during which players participated in an average of 45.5 official sets. For each player, we recorded three individual-level variables used in the following analyses: (1) Position, the on-court role (middle blocker, setter, libero, opposite hitter, and outside hitter) based on official match rosters, and all players had trained in their current position for at least four years; (2) Sets, the number of sets played during the season; and (3) Scores, the total points scored during the season.

The study was approved by the NYCU Research Ethics Center (YM111037e), and all participants gave written informed consent before participation.

### Stimuli

Participants viewed a 9 min 47 s compilation of highlights from three 2022 FIVB Volleyball Men’s Nations League matches (China vs. Japan, Japan vs. Italy, and Poland vs. USA), presented via rear projection and mirror. To create the experimental stimulus, replay scenes, close-up shots, and commentary segments were removed, retaining only the gameplay sequences from the moment of serve to the end of each rally. This editing preserved uninterrupted volleyball play and reduced non-game-related content.

The clips were selected to meet three criteria adapted from Parkinson et al. (2018): (1) the matches were unlikely to have been previously seen by participants, thereby reducing potential memory or familiarity biases; (2) the high-level international gameplay was likely to sustain attentional involvement; and (3) the clips were intended to provide a dynamic context in which individual differences in attention, emotional appraisal, and interpretation could naturally emerge during viewing.

### Imaging Acquisition

Participants were scanned at National Yang Ming Chiao Tung University using a 3T Siemens Trio scanner with a 32-channel head coil. A T1-weighted anatomical image was acquired using an MPRAGE sequence (TR = 2530 ms, TE = 3.03 ms, field of view = 220 mm, voxel size = 1×1×1 mm³). Functional images for both the game-viewing and resting-state runs were acquired using an echo-planar imaging sequence (TR = 2000 ms, TE = 30 ms, field of view = 220 mm, voxel size = 3.4×3.4×3.0 mm³). Inline retrospective motion correction was applied at the scanner during functional image acquisition (Lanka & Deshpande, 2019). The game-viewing run comprised 296 volumes (∼9 min 52 s) and was followed by the resting-state run, which comprised 180 volumes (∼6 min 08 s). During the resting-state run, participants were instructed to keep their eyes open, fixate on a central crosshair, avoid engaging in structured thoughts, and remain awake.

### Preprocessing

Data were preprocessed in CONN v22 and SPM12 under MATLAB R2023b following the default CONN pipeline (Nieto-Castanon, 2020; Whitfield-Gabrieli & Nieto-Castanon, 2012). Briefly, functional images were first realigned and unwrapped, slice-time corrected, and screened for ART outliers (framewise displacement > 0.9 mm or global BOLD signal changes > 5 standard deviations). Structural images were segmented into gray matter, white matter, and cerebrospinal fluid, and both structural and functional images were normalized to Montreal Neurological Institute (MNI) space. Functional images were then smoothed with an 8 mm full-width at half-maximum Gaussian kernel. For denoising, confounding effects were regressed out, including six motion parameters and their derivatives, anatomical component-based noise components from white matter and cerebrospinal fluid, ART-identified outlier scans, and linear trends. A band-pass filter (0.008–0.09 Hz) was then applied.

### Functional Connectivity

To examine functional connectivity at the level of large-scale brain networks, we used the Schaefer et al. (2018) cortical parcellation, which divides the cortex into 100 parcels assigned to one of seven networks based on Yeo et al. (2011). Each parcel was treated as a region of interest (ROI). For each participant, a 100 × 100 functional connectivity matrix was constructed by computing bivariate Pearson’s correlations between the mean BOLD time series of all possible pairs of parcels. Correlation coefficients were Fisher z-transformed before analysis. The 4,950 unique ROI-to-ROI connectivity values from the upper triangle of the matrix, excluding the diagonal, were used to represent each participant’s whole-brain functional connectivity profile.

### Intersubject differences

#### Neural differences

We quantified inter-participant differences in network-level functional connectivity in two steps (**Figure 1**). First, for each condition (resting-state and game-viewing), the 4,950 ROI-to-ROI functional connectivity values were aggregated into within- and between-network measures. Within-network functional connectivity was calculated as the mean of all connectivity values between ROIs within the same network, yielding seven values (one per network). Similarly, between-network functional connectivity was calculated as the mean of all connectivity values between ROIs from different networks, yielding 21 values corresponding to the unique network pairs. This procedure resulted in a 28-value network-level functional connectivity profile for each participant in each condition. Second, for each pair of participants, we computed the absolute difference between their network-level functional connectivity values.

**Figure 1.**
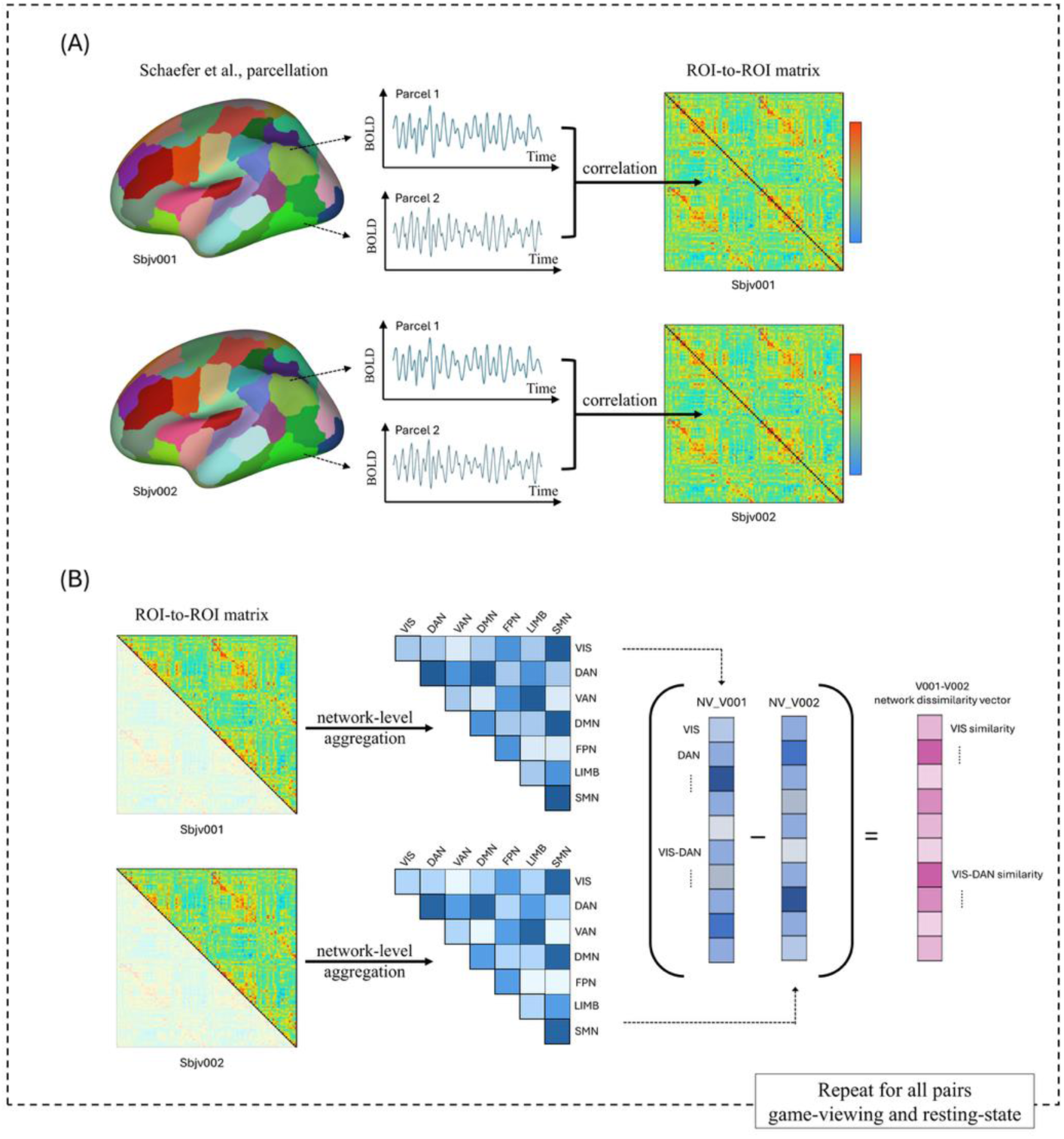
Computation of intersubject differences in functional connectivity. (A) The Schaefer et al. (2018) atlas was used to identify 100 cortical parcels assigned to one of seven networks defined by Yeo et al. (2011). For each participant, a 100 × 100 functional connectivity matrix was constructed by computing bivariate Pearson’s correlations between the mean BOLD time series of all possible ROI pairs. (B) For each participant, the 4,950 unique ROI-to-ROI connectivity values from the upper triangle of the functional connectivity matrix, excluding the diagonal. The ROI-level matrix was then summarized into 28 network-level FC measures. Intersubject differences in functional connectivity between a pair of participants were then computed as the absolute differences between their respective network-level profiles.

#### Non-neural differences

Position difference was coded as 1 when two players occupied the same position and 2 otherwise. Similarly, Team difference was coded as 1 when two players belonged to the same team and 2 otherwise. We coded the quantitative difference as 2 and 1 instead of 1 and 0, as the former way allows us to construct interaction terms of the variables to be assessed in regression model. To account for potential confounding factors that might covary with court position, we additionally included two player-level variables, Sets and Scores. Dyadic differences in Sets and Scores were calculated as the absolute differences between the two players’ values.

### Linear mixed-effects models

To test whether inter-participant differences in network-level functional connectivity were associated with players’ court position and team membership, we employed linear mixed-effects models using the R packages *lme4* and *lmerTest*. The unit of analysis was the dyad (i.e., a pair of participants). To account for the non-independence introduced by repeated observations of the same participants across dyads, we made the following adjustments in reference to Chen et al. (2017) and Hyon, Youm, et al. (2020). We duplicated the dyadic dataset by reversing the order of participants within each pair, which allowed us to specify crossed random effects for the two members of each dyad. Standard errors were adjusted by a scaling factor of 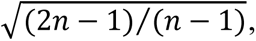, where *n* denotes the number of participants. Degrees of freedom for the *t* test were adjusted from 2*N*−*k*−1 to *N*−*k*−1, where *k* is the number of fixed-effect variables and *N* is the number of dyads in the undoubled dataset. *P*-values were further corrected for multiple comparisons across networks using the Benjamini–Hochberg false discovery rate (FDR) procedure.

These analyses were conducted separately for each of the 28 network-level functional connectivity measures (7 within-network and 21 between-network) in both resting-state and game-viewing conditions. For each network measure and condition, we first fitted a model including Position, Team, and their interaction, followed by a model that additionally included the control variables Sets and Scores. As a supplementary analysis to further evaluate the contribution of these control variables, we also fitted additional models in which Sets or Scores were treated as the primary variables of interest.

We assessed collinearity using generalized variance inflation factors (GVIF; Fox and Monette (1992), which are better suited than conventional VIF for models including interaction terms and non-independent observations. Because interaction terms can introduce inevitably inflated collinearity, we followed the sum-to-zero (effect) coding approach proposed by Kim and Jung (2025) to reduce non-essential collinearity between main effects and interaction terms. For this diagnostic only, categorical variables were effect-coded such that the coefficients across levels summed to zero. The GVIF diagnostics shows no warnings of collinearity (see **Table S1**).

### Machine learning classification

As a complementary analysis based on high-dimensional dyadic connectivity differences, we trained a machine learning classifier to predict whether two players occupied the same court position from inter-participant differences in whole-brain functional connectivity. For each participant pair, the input consisted of the absolute differences between corresponding 4,950 ROI-to-ROI connectivity values from the upper triangle of the two participants’ functional connectivity matrices, excluding the diagonal. The output of the model was a binary label indicating whether the two players occupied the same position. With 25 players, this resulted in 300 unique participant pairs. Classification analyses were conducted separately for the resting-state and game-viewing conditions. For each condition, we evaluated two input feature configurations: (1) functional connectivity differences only and (2) functional connectivity differences combined with Team label. To prevent data leakage, data splitting was performed at the participant level rather than at the dyad level. This procedure ensured that the model learned patterns of intersubject differences rather than participant-specific features. We employed *k*-fold cross-validation (*k* = 5) to evaluate model performance and summarized using accuracy, F1 score, recall, and precision. As a baseline, we also evaluated a frequency-based classifier, i.e., prediction based on the proportion of same-and different-position dyads in the data.

Several additional steps were implemented to improve model training. First, dimensionality reduction was performed using principal component analysis. The optimal number of dimensions was selected using the elbow method based on the relationship between model performance and dimensionality. Second, because the dataset was imbalanced, with unequal numbers of players across positions, we addressed class imbalance using synthetic data generation. Specifically, for each sample in the minority class, the algorithm identified its five nearest neighbors, based on Euclidean distance, within the same class. Synthetic samples were then generated by interpolating between the original sample and its selected neighbors. Finally, we implemented a multilayer perceptron (MLP) as the classification model, consisting of two hidden layers with 128 units each and a dropout rate of 0.5.

## Results

### Descriptive statistics of intersubject differences

A total of 300 unique dyads were generated from the 25 players, including 150 within-team and 150 between-team dyads. Of these dyads, 62 were same-position dyads and 238 were different-position dyads. Across all dyads, the mean difference in Sets was 12.4 (SD = 8.8) and the mean difference in Scores was 50.4 points (SD = 48.5) (**Table 2**).

**Table 2.** Dyadic differences in player characteristics within and between teams.

|  | All dyads<br>(n = 300) | Within team dyads<br>(n = 150) | Between team dyads<br>(n = 150) |
| --- | --- | --- | --- |
| Same-position dyads (n) | 62 | 28 | 34 |
| Sets difference (sets) | 12.4 ± 8.8 | 12.2 ± 8.7 | 12.6 ± 8.9 |
| Scores difference (points) | 50.4 ± 48.5 | 53.1 ± 51.7 | 47.6 ± 45.0 |

### Results of linear mixed-effects models

Using linear mixed-effects models, we estimated regression coefficients (*β*) and FDR-corrected *p* values for the resting-state and game-viewing conditions. In the resting-state condition, statistically significant results were only found within the somatomotor network (SMN), as detailed below. On the other hand, in the game-viewing condition, significant effects were observed across many network pairs. Robust effects (FDR-corrected *p* < 0.001) are frequently observed in network pairs involving the default mode network (DMN), SMN, and frontoparietal network, especially when interacting with the visual network. In the models without covariates, 15 of 28 network pairs showed a coherent pattern of Position and Team main effects, with negative Position × Team interactions. After including Sets and Scores as covariates, significant main effects of Position and Team remained evident across 12 of 28 network pairs, while the Position × Team interaction became limited. Because the game-viewing results were widespread, **Table 3** highlights the key network pairs showing significant Position × Team interactions in the models with control variables; broader visualization and full results are provided in **Figure 2** and **Table S2**, respectively.

**Figure 2.**
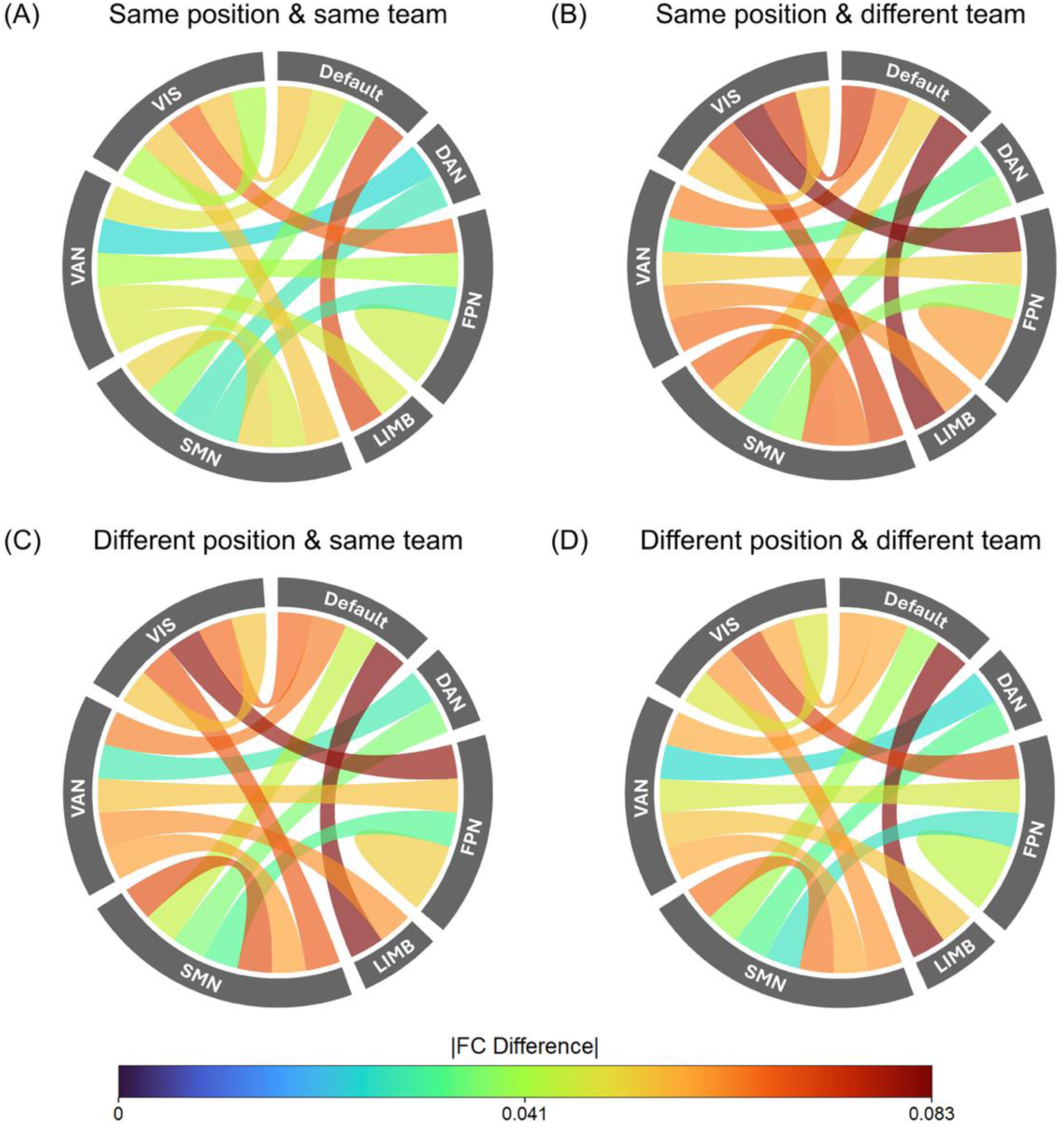
Significant inter-participant differences in network-level functional connectivity in the game-viewing condition. Chord diagrams visualize the mean absolute differences in network-level functional connectivity across participant pairs for network measures showing significant effects in the linear mixed-effects models. Ribbon color indicates the magnitude of the average absolute functional connectivity difference, with cooler colors indicating smaller differences and warmer colors indicating larger differences. The visualization is divided into four subsets based on whether two players shared the same court position and whether they belonged to the same team: (A) same position and same team, (B) same position and different team, (C) different position and same team, and (D) different position and different team. *Notes*. DAN: Dorsal attention network, Default: Default mode network, FPN: Frontoparietal network, LIMB: Limbic network, SMN: Somatomotor network, VAN: Salience/Ventral attention network, VIS: Visual network.

**Table 3.**
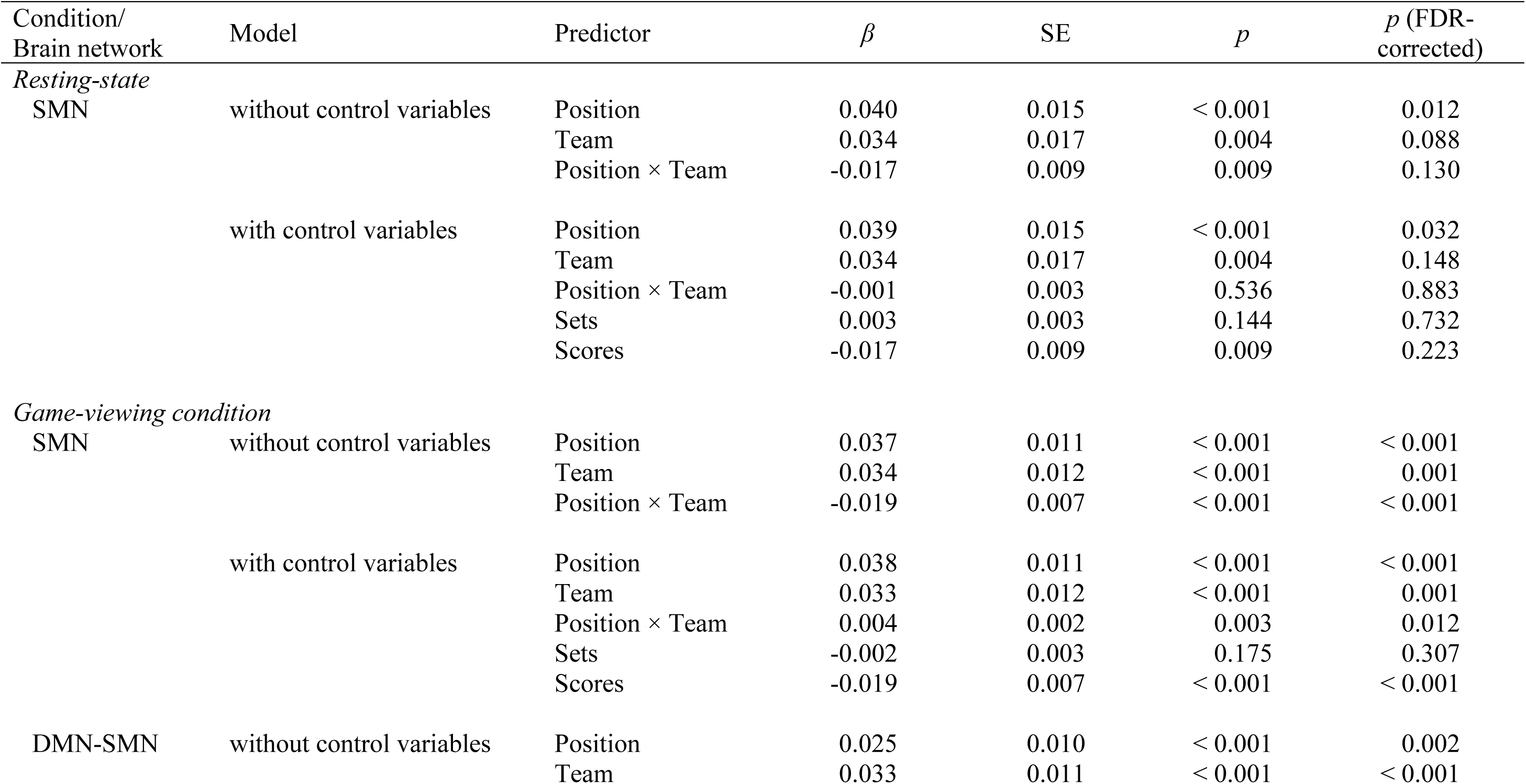
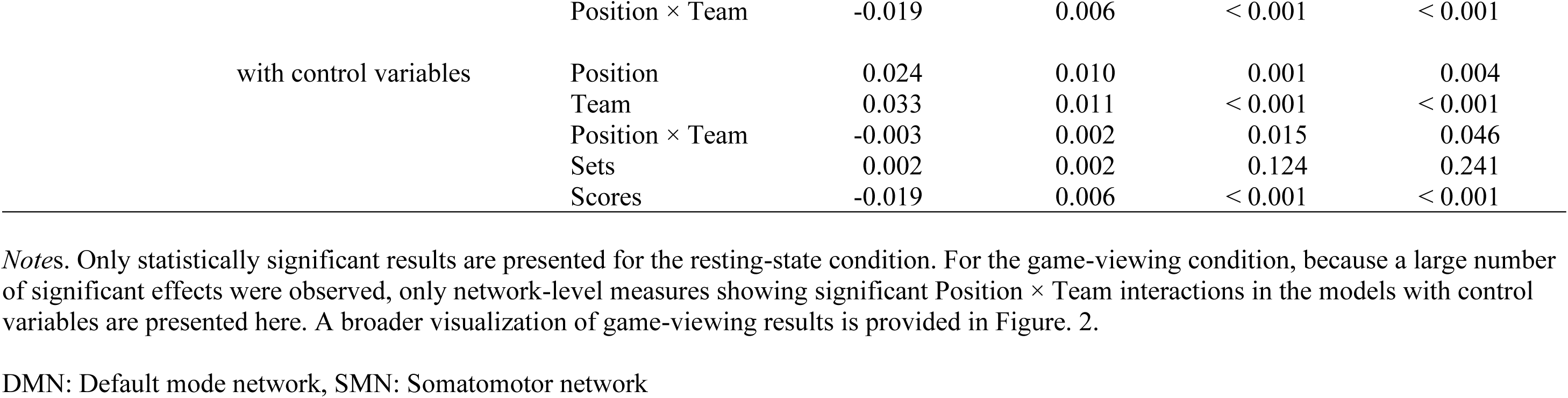
Results of linear mixed-effects models predicting inter-participant differences in network-level functional connectivity from Position and Team, with and without control variables.

In the resting-state condition, only functional connectivity differences within the SMN were significantly associated with Position (*β* = 0.040, FDR-corrected *p* = 0.012), indicating that dyads sharing the same position exhibited smaller SMN functional connectivity differences. The main effects of Team and the Position × Team interaction were not significant after FDR correction. With covariate adjustment, the Position effect in the SMN remained significant (*β* = 0.039, FDR-corrected *p* = 0.032), whereas the other effects did not.

In the game-viewing condition, functional connectivity differences were associated with Position, Team, and their interaction across multiple network-level measures (see **Figure 2** for illustration and **Table S2** for details). In the models without covariates, both the SMN and the connectivity between the DMN and SMN showed positive main effects of Position and Team, together with negative Position × Team interactions (SMN: Position *β* = 0.037, FDR-corrected *p* < 0.001; Team *β* = 0.034, FDR-corrected *p* = 0.001; Position × Team *β* = -0.019, FDR-corrected *p* < 0.001; DMN–SMN: Position *β* = 0.025, FDR-corrected *p* = 0.002; Team *β* = 0.033, FDR-corrected *p* < 0.001; Position × Team *β* = -0.019, FDR-corrected *p* < 0.001). Given our coding, these positive main effects indicate smaller functional connectivity differences among dyads sharing the same position or team membership. The negative Position × Team interactions further indicate that the association between shared position and smaller functional connectivity differences was stronger for within-team dyads than for between-team dyads, consistent with our hypothesis. With covariate adjustment, the main effects of Position and Team remained significant in both network measures (SMN: Position *β* = 0.038, FDR-corrected *p* < 0.001; Team *β* = 0.033, FDR-corrected *p* = 0.001; DMN–SMN: Position *β* = 0.024, FDR-corrected *p* = 0.004; Team *β* = 0.033, FDR-corrected *p* < 0.001). However, the direction of the interaction varied across networks: the Position × Team term remained negative for DMN–SMN connectivity (*β* = -0.003, FDR-corrected *p* = 0.046), but was positive for within-SMN connectivity (*β* = 0.004, FDR-corrected *p* = 0.012). The positive Position × Team interaction within the SMN indicates that the association between shared position and smaller functional connectivity differences was stronger for between-team dyads than for within-team dyads, contrary to our hypothesis that team membership would increase the Position effect. Results from the supplementary models treating Sets or Scores as the primary variables of interest are reported in **Tables S3** and **S4** and are not discussed further here.

### Machine learning prediction

The MLP classifier predicted whether two players shared the same court position from differences in their functional connectivity patterns, achieving accuracies of 82% in the resting-state condition and 86% in the game-viewing condition (see **Table 4**). These prediction accuracies exceeded the frequency baseline accuracy of 79.33%. Notably, when team membership was added as an additional input alongside functional connectivity differences, classification accuracy further increased to 90% in the game-viewing condition. In contrast, adding team membership did not improve accuracy in the resting-state condition, which remained at 82%.

**Table 4.** Classification performance of the machine learning classifier for predicting whether two players shared the same court position.

| Metric | Resting-state condition |  | Game-viewing condition |  | Majority-class<br>baseline: Frequency |
| --- | --- | --- | --- | --- | --- |
|  | functional connectivity<br>data only | functional connectivity<br>data + Team | functional connectivity<br>data only | functional connectivity<br>data + Team |  |
| Accuracy | 82.00% | 82.00% | 86.00% | 90.00% | 79.33% |
| F1 Score | 82.72% | 82.72% | 83.23% | 85.26% | 70.18% |
| Recall Rate | 82.00% | 82.00% | 86.00% | 84.26% | 79.33% |
| Precision Rate | 83.48% | 83.48% | 80.63% | 88.00% | 62.93% |

## Discussion

In this study, we recruited professional volleyball players with long-term team training histories and stable court positions to examine whether cross-individual differences in functional connectivity are associated with dyadic similarity in position and team membership in a naturalistic context of team coordination. We found that players occupying the same position showed smaller functional connectivity differences, with these effects appearing most clearly during naturalistic game viewing. This position-related effect within the SMN was also observed during resting state. Shared team membership was also associated with smaller functional connectivity differences during game viewing, particularly in interaction with position, although the direction of this interaction varied across functional networks after controlling for Sets and Scores. These associations were further supported by a deep learning classifier showing that shared court position could be predicted from dyadic differences in functional connectivity and team membership with accuracy exceeding the frequency-based baseline. Prior work on functional connectome “fingerprinting” has emphasized the stability and individuality of brain network organization (Finn et al., 2015); in contrast, our findings suggest that such individual patterns may also be partially structured by shared role-specific and team-based experience.

More broadly, our findings extend prior work on neural similarity by identifying an organizational-level convergence within a real-world performance group. Whereas previous studies have linked neural similarity to social network closeness and shared cultural background (Hyon, Kleinbaum, et al., 2020; Hyon, Youm, et al., 2020; Luo et al., 2022), our results suggest that similar convergence may also emerge among individuals who occupy the same functional role within the same task-oriented team. Notably, the observed effects were more pronounced during naturalistic viewing than during the resting-state condition, suggesting that this form of neural similarity may be especially evident when participants engage with task-relevant stimuli that recruit learned position- or team-specific demands. In this line, our findings offer a neural perspective on ideas often discussed under the label of team chemistry, suggesting that coordinated group experience may be reflected in shared functional brain organization. These suggestions raise important questions for future research: How long does it take for neural similarity to form? Under what contexts does it emerge? And for which cognitive or functional processes? Addressing these questions will be an important direction for future work.

### Findings within somatomotor network

The SMN showed a robust association between shared position and smaller functional connectivity differences in both the resting-state and game-viewing conditions. Given that functional connectivity may reflect intrinsic network organization shaped by long-term experience and plasticity (Buckner et al., 2013), the SMN finding suggests that shared position-specific experience may contribute to convergent organization. Volleyball players who occupy the same position typically undergo similar motor training and repeatedly engage similar action-perception demands, making the SMN a plausible network in which role-specific experience would be expressed across both resting-state and naturalistic viewing conditions. The SMN is a robust intrinsic network encompassing primary motor and primary somatosensory cortices (Yeo et al., 2011). First, evidence that motor training can shape intrinsic sensorimotor connectivity comes from motor-learning studies, which showed functionally specific changes in resting-state sensorimotor connectivity following motor learning and experience (McGregor & Gribble, 2015; Vahdat et al., 2011). Second, evidence suggests that early somatosensory regions contribute to functions beyond action execution, including action observation and action-related perceptual inference (Valchev et al., 2016; Valchev et al., 2017). Specifically, perturbation of primary somatosensory cortex can alter resting-state functional connectivity within a broader somatosensory–motor circuit (Valchev et al., 2015), and action-perception processes during action observation are modulated by domain-specific perceptual and motor experience (Chen et al., 2023). Together, these findings are consistent with the possibility that same-position players develop partially convergent SMN organization through long-term role-specific training and action-perception experience. This interpretation is further supported by behavioral and neurophysiological evidence showing that action prediction improves when observed actions align with an athlete’s motor repertoire or position-specific expertise (Aglioti et al., 2008; Paolini et al., 2025). Notably, our static functional connectivity measure cannot directly index time-locked action-related processing during game viewing. Future studies could use methods such as intersubject correlation to further examine the mechanisms underlying position-specific similarity.

Beyond motor functions, the SMN has also been linked to socially relevant processes (Tompson et al., 2020; Volynets et al., 2020). For example, Volynets et al. (2020) reported high cross-modal classification accuracy for emotional facial expressions in primary motor and somatosensory cortices, suggesting that regions traditionally considered motor can encode socially relevant information. Likewise, Tompson et al. (2020) demonstrated that functional connectivity involving the DMN and SMN is altered during social network learning, indicating that social learning engages brain systems linked to action-related processing. This view may help contextualize the Team and Position main effects observed in the SMN during game viewing. Volleyball players sharing the same position typically have similar role-specific coordination patterns, including patterns of communication and repeated ball exchange confined by rotation rules and positional rules. Thus, smaller SMN connectivity differences may reflect not only shared motor repertoire but also socially structured role demands. Importantly, however, the *positive* Position × Team interaction in the covariates-adjusted SMN model indicates that the association between shared position and smaller SMN connectivity differences was not stronger for within-team dyads, which suggests that position-related SMN similarity may reflect a combination of shared motor repertoire and role-specific coordination experience that can extend across teams.

### *Negative* Position × Team interaction in default mode–somatomotor connectivity

In contrast to the within-SMN result, the covariates-adjusted DMN–SMN model showed a negative Position × Team interaction, indicating that shared position was associated with smaller connectivity differences more strongly among within-team than between-team dyads. This pattern suggests a more specific role of team membership in shaping position-related similarity in this network pair involving the DMN. The DMN is known to be highly individualized and idiosyncratic across individuals (Yeshurun et al., 2021), which means similarity in the DMN among individuals who share similar backgrounds may stand out in contrast to those who come from more divergent backgrounds. Moreover, DMN activity has been linked to predicting others’ behaviors during social interaction (Carter et al., 2012), and such predictive processing depends in part on how individuals interpret ongoing stimuli (Nguyen et al., 2019). In a naturalistic volleyball-viewing condition, players who occupy the same position within the same team may be more likely to interpret unfolding plays through similar role-specific and team-specific coordination experiences. From this perspective, greater similarity in DMN–SMN connectivity may reflect more similar coupling between higher-order interpretive processes and action-related areas during game viewing. This interpretation remains tentative, however, and future work combining advanced fMRI analyses with eye-tracking, behavioral judgments, and self-reports of event interpretation could help clarify whether the observed pattern reflects shared interpretation and meaning construction, or instead arises from shared attentional allocation.

### Position and Team effects during naturalistic game-viewing

The game-viewing condition showed widespread Position- and Team-related effects across network pairs, frequently including the DMN, SMN, frontoparietal, and visual networks. We suggest that long-term role-specific and team-based experience may shape how players process unfolding game scenes in this context. Prior work suggests that movie watching recruits not only sensory systems but also higher-order association systems supporting attention, executive control, and internally guided processing (Demirtas et al., 2019; Kim et al., 2018; Rajimehr et al., 2024). Specifically, movie watching has been shown to induce condition-specific functional reorganization involving particularly in frontal and parietal association regions (Demirtas et al., 2019). Other work suggests that naturalistic viewing increases interaction between systems associated with externally oriented perception and those associated with internally oriented control (Kim et al., 2018) and engages broad sensory, attentional, executive, language, and social-cognitive cortical systems (Rajimehr et al., 2024). In the context of volleyball, such a viewing state may allow long-term role-specific and team-based experience to shape how players allocate attention, perceive actions, and interpret unfolding play. Consistent with this possibility, athletes are known to deploy domain-specific attentional strategies during action observation (Meng et al., 2019). Accordingly, beyond the motor- and coordination-based accounts discussed above, our findings are consistent with the possibility that naturalistic volleyball scenes engage more similar modes of ongoing scene processing among players who share role-specific or team-based experience.

### Limitations

Our study has several limitations that warrant consideration in future research. First, we did not collect interpersonal or team-level behavioral measures that could help us to directly explain social interaction and coordination among players, such as social network structure, perceived team cohesion, team identification, or shared tactical understanding. Those measures would be valuable for clarifying how team culture, social dynamics, and training experiences are associated with similarity in functional connectivity. Second, because our data were cross-sectional, we cannot draw strong causal conclusions. The observed associations may reflect the effects of long-term shared training and team experience, but they may also partly reflect self-selection, in which individuals with particular traits are more likely to occupy certain positions or remain in specific team environments. Longitudinal designs with repeated neuroimaging across training periods would be better suited to disentangling experience-dependent change from pre-existing individual differences, and to determining whether and how prolonged role and team training shapes similarity in functional connectivity over time. Finally, the generalizability of the present findings is limited by modest sample size and specific sample composition. Although the dyadic design yielded 300 participant pairs, the study included only 25 male elite volleyball players from two teams. Future research with larger and more diverse samples will be needed to determine whether similar patterns are observed across other teams, competitive levels, sports, and populations.

## Acknowledgments

We thank Jin Liu for assistance with data cleaning and Hsiang-Hisen Chang for assistance with implementing the machine learning model

## Data availability statement

The data underlying this article will be shared on reasonable request to the corresponding author.

## AI assistance disclosure

We disclose the use of AI-assisted tools during manuscript preparation. ChatGPT and Quillbot were used to assist with readability improvement and grammar checking. The authors reviewed and verified all AI-assisted outputs and take full responsibility for the content of the manuscript.

## Author contributions

J.J.C.: Conceptualization (Equal), Formal analysis (Equal), Visualization (Lead), Writing—original draft (Equal).

Y.C.C.: Conceptualization (Equal), Funding acquisition (Equal), Investigation (Lead), Supervision (Equal), Writing—review and editing (Supporting).

Y.S.C.: Conceptualization (Equal), Formal analysis (Equal), Visualization (Supporting), Funding acquisition (Equal); Supervision (Equal), Writing—original draft (Equal); Writing—review and editing (Lead).

