## Supplemental information for "Shared roles and team membership are reflected in functional connectome similarity: Neural evidence from real-world volleyball teams"

### Supplementary

**Table S1-1** GVIF-based collinearity diagnostics for the Position × Team models with control variables in the resting-state condition.

| Brain network(s) | Variables | GVIF | VIF 95% CI lower | VIF 95% CI upper | adjusted VIF |
|---|---|---|---|---|---|
| DAN-DAN | Team | 1.087 | 1.020 | 1.383 | 1.043 |
| | Position | 1.064 | 1.009 | 1.445 | 1.031 |
| | Sets | 1.099 | 1.026 | 1.375 | 1.048 |
| | Scores | 1.144 | 1.054 | 1.385 | 1.070 |
| | Position × Team | 1.096 | 1.024 | 1.376 | 1.047 |
| DAN-DMN | Team | 1.104 | 1.029 | 1.373 | 1.051 |
| | Position | 1.063 | 1.009 | 1.452 | 1.031 |
| | Sets | 1.100 | 1.027 | 1.374 | 1.049 |
| | Scores | 1.145 | 1.054 | 1.386 | 1.070 |
| | Position × Team | 1.111 | 1.033 | 1.372 | 1.054 |
| DAN-FPN | Team | 1.098 | 1.025 | 1.375 | 1.048 |
| | Position | 1.063 | 1.009 | 1.452 | 1.031 |
| | Sets | 1.101 | 1.027 | 1.374 | 1.049 |
| | Scores | 1.146 | 1.055 | 1.386 | 1.070 |
| | Position × Team | 1.106 | 1.030 | 1.373 | 1.052 |
| DAN-LIMB | Team | 1.104 | 1.029 | 1.373 | 1.051 |
| | Position | 1.063 | 1.009 | 1.452 | 1.031 |
| | Sets | 1.100 | 1.027 | 1.374 | 1.049 |
| | Scores | 1.145 | 1.054 | 1.386 | 1.070 |
| | Position × Team | 1.111 | 1.033 | 1.372 | 1.054 |
| DAN-SMN | Team | 1.051 | 1.005 | 1.552 | 1.025 |

| | | | | | |
|---|---|---|---|---|---|
| | Position | 1.067 | 1.010 | 1.433 | 1.033 |
| | Sets | 1.097 | 1.025 | 1.376 | 1.047 |
| | Scores | 1.143 | 1.053 | 1.385 | 1.069 |
| | Position × Team | 1.061 | 1.008 | 1.460 | 1.030 |
| DAN-VAN | Team | 1.100 | 1.027 | 1.374 | 1.049 |
| | Position | 1.063 | 1.009 | 1.450 | 1.031 |
| | Sets | 1.099 | 1.026 | 1.375 | 1.048 |
| | Scores | 1.145 | 1.054 | 1.385 | 1.070 |
| | Position × Team | 1.108 | 1.031 | 1.373 | 1.052 |
| DAN-VIS | Team | 1.043 | 1.003 | 1.684 | 1.021 |
| | Position | 1.067 | 1.010 | 1.431 | 1.033 |
| | Sets | 1.097 | 1.025 | 1.376 | 1.047 |
| | Scores | 1.143 | 1.053 | 1.385 | 1.069 |
| | Position × Team | 1.054 | 1.006 | 1.512 | 1.027 |
| DMN-DMN | Team | 1.113 | 1.034 | 1.373 | 1.055 |
| | Position | 1.064 | 1.009 | 1.444 | 1.032 |
| | Sets | 1.095 | 1.024 | 1.377 | 1.046 |
| | Scores | 1.144 | 1.054 | 1.385 | 1.069 |
| | Position × Team | 1.117 | 1.037 | 1.373 | 1.057 |
| DMN-FPN | Team | 1.066 | 1.010 | 1.434 | 1.033 |
| | Position | 1.066 | 1.010 | 1.436 | 1.032 |
| | Sets | 1.096 | 1.025 | 1.376 | 1.047 |
| | Scores | 1.143 | 1.053 | 1.385 | 1.069 |

| | | | | | |
|---|---|---|---|---|---|
| | Position × Team | 1.075 | 1.014 | 1.404 | 1.037 |
| DMN-LIMB | Team | 1.090 | 1.021 | 1.381 | 1.044 |
| | Position | 1.064 | 1.009 | 1.445 | 1.031 |
| | Sets | 1.098 | 1.026 | 1.375 | 1.048 |
| | Scores | 1.144 | 1.054 | 1.385 | 1.070 |
| | Position × Team | 1.098 | 1.025 | 1.375 | 1.048 |
| DMN-SMN | Team | 1.062 | 1.009 | 1.453 | 1.031 |
| | Position | 1.066 | 1.010 | 1.436 | 1.032 |
| | Sets | 1.097 | 1.025 | 1.376 | 1.047 |
| | Scores | 1.143 | 1.053 | 1.385 | 1.069 |
| | Position × Team | 1.072 | 1.013 | 1.413 | 1.035 |
| DMN-VAN | Team | 1.059 | 1.007 | 1.476 | 1.029 |
| | Position | 1.066 | 1.010 | 1.434 | 1.033 |
| | Sets | 1.097 | 1.025 | 1.376 | 1.047 |
| | Scores | 1.143 | 1.053 | 1.385 | 1.069 |
| | Position × Team | 1.068 | 1.011 | 1.425 | 1.034 |
| DMN-VIS | Team | 1.053 | 1.005 | 1.523 | 1.026 |
| | Position | 1.066 | 1.010 | 1.434 | 1.033 |
| | Sets | 1.097 | 1.025 | 1.375 | 1.047 |
| | Scores | 1.143 | 1.053 | 1.385 | 1.069 |
| | Position × Team | 1.064 | 1.009 | 1.446 | 1.031 |
| FPN-FPN | Team | 1.080 | 1.016 | 1.394 | 1.039 |
| | Position | 1.064 | 1.009 | 1.445 | 1.031 |

| | | | | | |
|---|---|---|---|---|---|
| | Sets | 1.100 | 1.027 | 1.374 | 1.049 |
| | Scores | 1.145 | 1.054 | 1.386 | 1.070 |
| | Position × Team | 1.089 | 1.021 | 1.381 | 1.044 |
| FPN-LIMB | Team | 1.091 | 1.022 | 1.380 | 1.045 |
| | Position | 1.063 | 1.009 | 1.448 | 1.031 |
| | Sets | 1.099 | 1.026 | 1.374 | 1.048 |
| | Scores | 1.145 | 1.054 | 1.386 | 1.070 |
| | Position × Team | 1.099 | 1.026 | 1.374 | 1.049 |
| FPN-SMN | Team | 1.028 | 1.000 | 2.698 | 1.014 |
| | Position | 1.068 | 1.011 | 1.426 | 1.034 |
| | Sets | 1.096 | 1.024 | 1.376 | 1.047 |
| | Scores | 1.142 | 1.053 | 1.384 | 1.069 |
| | Position × Team | 1.040 | 1.002 | 1.767 | 1.020 |
| FPN-VAN | Team | 1.067 | 1.010 | 1.433 | 1.033 |
| | Position | 1.065 | 1.010 | 1.439 | 1.032 |
| | Sets | 1.098 | 1.026 | 1.375 | 1.048 |
| | Scores | 1.144 | 1.054 | 1.385 | 1.070 |
| | Position × Team | 1.076 | 1.014 | 1.402 | 1.037 |
| FPN-VIS | Team | 1.077 | 1.015 | 1.400 | 1.038 |
| | Position | 1.064 | 1.009 | 1.444 | 1.032 |
| | Sets | 1.099 | 1.026 | 1.374 | 1.048 |
| | Scores | 1.145 | 1.054 | 1.385 | 1.070 |
| | Position × Team | 1.086 | 1.019 | 1.384 | 1.042 |

| | | | | | |
|---|---|---|---|---|---|
| LIMB-LIMB | Team | 1.173 | 1.074 | 1.407 | 1.083 |
| | Position | 1.058 | 1.007 | 1.484 | 1.028 |
| | Sets | 1.105 | 1.030 | 1.373 | 1.051 |
| | Scores | 1.148 | 1.057 | 1.388 | 1.072 |
| | Position × Team | 1.177 | 1.077 | 1.411 | 1.085 |
| LIMB-SMN | Team | 1.106 | 1.030 | 1.373 | 1.051 |
| | Position | 1.062 | 1.008 | 1.454 | 1.031 |
| | Sets | 1.101 | 1.027 | 1.374 | 1.049 |
| | Scores | 1.146 | 1.055 | 1.386 | 1.070 |
| | Position × Team | 1.113 | 1.034 | 1.373 | 1.055 |
| LIMB-VAN | Team | 1.094 | 1.024 | 1.377 | 1.046 |
| | Position | 1.064 | 1.009 | 1.447 | 1.031 |
| | Sets | 1.098 | 1.026 | 1.375 | 1.048 |
| | Scores | 1.144 | 1.054 | 1.385 | 1.070 |
| | Position × Team | 1.102 | 1.028 | 1.374 | 1.050 |
| LIMB-VIS | Team | 1.046 | 1.003 | 1.635 | 1.023 |
| | Position | 1.067 | 1.010 | 1.432 | 1.033 |
| | Sets | 1.097 | 1.025 | 1.375 | 1.047 |
| | Scores | 1.143 | 1.053 | 1.385 | 1.069 |
| | Position × Team | 1.057 | 1.006 | 1.493 | 1.028 |
| SMN-SMN | Team | 1.250 | 1.129 | 1.483 | 1.118 |
| | Position | 1.060 | 1.008 | 1.465 | 1.030 |
| | Sets | 1.093 | 1.023 | 1.378 | 1.045 |

| | | | | | |
|---|---|---|---|---|---|
| | Scores | 1.143 | 1.053 | 1.384 | 1.069 |
| | Position × Team | 1.246 | 1.126 | 1.479 | 1.116 |
| SMN-VAN | Team | 1.059 | 1.007 | 1.477 | 1.029 |
| | Position | 1.066 | 1.010 | 1.436 | 1.032 |
| | Sets | 1.098 | 1.026 | 1.375 | 1.048 |
| | Scores | 1.144 | 1.054 | 1.385 | 1.069 |
| | Position × Team | 1.069 | 1.011 | 1.424 | 1.034 |
| SMN-VIS | Team | 1.080 | 1.016 | 1.394 | 1.039 |
| | Position | 1.065 | 1.010 | 1.439 | 1.032 |
| | Sets | 1.096 | 1.025 | 1.376 | 1.047 |
| | Scores | 1.144 | 1.054 | 1.385 | 1.069 |
| | Position × Team | 1.088 | 1.020 | 1.383 | 1.043 |
| VAN-VAN | Team | 1.063 | 1.009 | 1.450 | 1.031 |
| | Position | 1.065 | 1.010 | 1.438 | 1.032 |
| | Sets | 1.098 | 1.026 | 1.375 | 1.048 |
| | Scores | 1.144 | 1.054 | 1.385 | 1.070 |
| | Position × Team | 1.073 | 1.013 | 1.410 | 1.036 |
| VAN-VIS | Team | 1.051 | 1.005 | 1.552 | 1.025 |
| | Position | 1.066 | 1.010 | 1.435 | 1.032 |
| | Sets | 1.098 | 1.026 | 1.375 | 1.048 |
| | Scores | 1.144 | 1.054 | 1.385 | 1.070 |
| | Position × Team | 1.062 | 1.008 | 1.458 | 1.030 |
| VIS-VIS | Team | 1.045 | 1.003 | 1.644 | 1.022 |

| | | | | | |
|---|---|---|---|---|---|
| | Position | 1.067 | 1.010 | 1.432 | 1.033 |
| | Sets | 1.097 | 1.025 | 1.375 | 1.048 |
| | Scores | 1.143 | 1.053 | 1.385 | 1.069 |
| | Position × Team | 1.056 | 1.006 | 1.496 | 1.028 |

**Table S1-2** GVIF-based collinearity diagnostics for the Position × Team models with control variables in the game-viewing condition.

| Brain network(s) | Variables | GVIF | VIF 95% CI lower | VIF 95% CI upper | adjusted VIF |
|---|---|---|---|---|---|
| DAN-DAN | Team | 1.111 | 1.033 | 1.372 | 1.054 |
| | Position | 1.062 | 1.009 | 1.454 | 1.031 |
| | Sets | 1.100 | 1.027 | 1.374 | 1.049 |
| | Scores | 1.145 | 1.055 | 1.386 | 1.070 |
| | Position × Team | 1.118 | 1.037 | 1.373 | 1.057 |
| DAN-DMN | Team | 1.085 | 1.019 | 1.386 | 1.042 |
| | Position | 1.063 | 1.009 | 1.447 | 1.031 |
| | Sets | 1.100 | 1.027 | 1.374 | 1.049 |
| | Scores | 1.145 | 1.055 | 1.386 | 1.070 |
| | Position × Team | 1.094 | 1.023 | 1.377 | 1.046 |
| DAN-FPN | Team | 1.068 | 1.011 | 1.426 | 1.034 |
| | Position | 1.065 | 1.010 | 1.440 | 1.032 |
| | Sets | 1.099 | 1.026 | 1.375 | 1.048 |
| | Scores | 1.144 | 1.054 | 1.385 | 1.070 |
| | Position × Team | 1.078 | 1.015 | 1.398 | 1.038 |

| | | | | | |
|---|---|---|---|---|---|
| DAN-LIMB | Team | 1.100 | 1.026 | 1.374 | 1.049 |
| | Position | 1.063 | 1.009 | 1.451 | 1.031 |
| | Sets | 1.100 | 1.027 | 1.374 | 1.049 |
| | Scores | 1.145 | 1.055 | 1.386 | 1.070 |
| | Position × Team | 1.107 | 1.031 | 1.373 | 1.052 |
| DAN-SMN | Team | 1.043 | 1.003 | 1.684 | 1.021 |
| | Position | 1.067 | 1.010 | 1.431 | 1.033 |
| | Sets | 1.097 | 1.025 | 1.376 | 1.047 |
| | Scores | 1.143 | 1.053 | 1.385 | 1.069 |
| | Position × Team | 1.054 | 1.006 | 1.512 | 1.027 |
| DAN-VAN | Team | 1.121 | 1.039 | 1.374 | 1.059 |
| | Position | 1.060 | 1.008 | 1.467 | 1.030 |
| | Sets | 1.105 | 1.030 | 1.373 | 1.051 |
| | Scores | 1.148 | 1.057 | 1.388 | 1.071 |
| | Position × Team | 1.128 | 1.043 | 1.376 | 1.062 |
| DAN-VIS | Team | 1.116 | 1.036 | 1.373 | 1.056 |
| | Position | 1.062 | 1.008 | 1.458 | 1.030 |
| | Sets | 1.101 | 1.027 | 1.374 | 1.049 |
| | Scores | 1.146 | 1.055 | 1.386 | 1.070 |
| | Position × Team | 1.123 | 1.040 | 1.375 | 1.060 |
| DMN-DMN | Team | 1.166 | 1.069 | 1.401 | 1.080 |
| | Position | 1.059 | 1.007 | 1.474 | 1.029 |
| | Sets | 1.102 | 1.028 | 1.374 | 1.050 |

| | | | | | |
|---|---|---|---|---|---|
| | Scores | 1.146 | 1.055 | 1.387 | 1.071 |
| | Position × Team | 1.170 | 1.072 | 1.405 | 1.082 |
| DMN-FPN | Team | 1.230 | 1.115 | 1.462 | 1.109 |
| | Position | 1.056 | 1.006 | 1.501 | 1.027 |
| | Sets | 1.105 | 1.030 | 1.373 | 1.051 |
| | Scores | 1.149 | 1.057 | 1.388 | 1.072 |
| | Position × Team | 1.232 | 1.116 | 1.464 | 1.110 |
| DMN-LIMB | Team | 1.172 | 1.073 | 1.406 | 1.083 |
| | Position | 1.059 | 1.007 | 1.475 | 1.029 |
| | Sets | 1.102 | 1.028 | 1.374 | 1.050 |
| | Scores | 1.146 | 1.055 | 1.387 | 1.071 |
| | Position × Team | 1.176 | 1.076 | 1.410 | 1.084 |
| DMN-SMN | Team | 1.129 | 1.044 | 1.377 | 1.063 |
| | Position | 1.060 | 1.008 | 1.469 | 1.029 |
| | Sets | 1.105 | 1.029 | 1.373 | 1.051 |
| | Scores | 1.148 | 1.056 | 1.388 | 1.071 |
| | Position × Team | 1.136 | 1.048 | 1.380 | 1.066 |
| DMN-VAN | Team | 1.160 | 1.065 | 1.397 | 1.077 |
| | Position | 1.058 | 1.007 | 1.482 | 1.029 |
| | Sets | 1.106 | 1.030 | 1.373 | 1.052 |
| | Scores | 1.149 | 1.057 | 1.388 | 1.072 |
| | Position × Team | 1.165 | 1.068 | 1.401 | 1.079 |
| DMN-VIS | Team | 1.154 | 1.060 | 1.392 | 1.074 |

| | | | | | |
|---|---|---|---|---|---|
| | Position | 1.058 | 1.007 | 1.478 | 1.029 |
| | Sets | 1.106 | 1.030 | 1.373 | 1.051 |
| | Scores | 1.148 | 1.057 | 1.388 | 1.072 |
| | Position × Team | 1.159 | 1.064 | 1.396 | 1.077 |
| FPN-FPN | Team | 1.404 | 1.247 | 1.661 | 1.185 |
| | Position | 1.054 | 1.006 | 1.511 | 1.027 |
| | Sets | 1.092 | 1.022 | 1.379 | 1.045 |
| | Scores | 1.140 | 1.051 | 1.383 | 1.068 |
| | Position × Team | 1.400 | 1.244 | 1.656 | 1.183 |
| FPN-LIMB | Team | 1.231 | 1.115 | 1.463 | 1.109 |
| | Position | 1.055 | 1.006 | 1.504 | 1.027 |
| | Sets | 1.106 | 1.030 | 1.373 | 1.052 |
| | Scores | 1.149 | 1.057 | 1.388 | 1.072 |
| | Position × Team | 1.233 | 1.116 | 1.465 | 1.110 |
| FPN-SMN | Team | 1.067 | 1.010 | 1.431 | 1.033 |
| | Position | 1.065 | 1.010 | 1.441 | 1.032 |
| | Sets | 1.099 | 1.026 | 1.374 | 1.048 |
| | Scores | 1.145 | 1.054 | 1.385 | 1.070 |
| | Position × Team | 1.077 | 1.015 | 1.401 | 1.038 |
| FPN-VAN | Team | 1.144 | 1.054 | 1.385 | 1.069 |
| | Position | 1.058 | 1.007 | 1.484 | 1.028 |
| | Sets | 1.110 | 1.032 | 1.372 | 1.053 |
| | Scores | 1.151 | 1.059 | 1.390 | 1.073 |

| | | | | | |
|---|---|---|---|---|---|
| | Position × Team | 1.149 | 1.057 | 1.389 | 1.072 |
| FPN-VIS | Team | 1.109 | 1.032 | 1.372 | 1.053 |
| | Position | 1.061 | 1.008 | 1.458 | 1.030 |
| | Sets | 1.103 | 1.028 | 1.373 | 1.050 |
| | Scores | 1.146 | 1.055 | 1.387 | 1.071 |
| | Position × Team | 1.116 | 1.036 | 1.373 | 1.057 |
| LIMB-LIMB | Team | 1.117 | 1.037 | 1.373 | 1.057 |
| | Position | 1.062 | 1.008 | 1.456 | 1.030 |
| | Sets | 1.100 | 1.027 | 1.374 | 1.049 |
| | Scores | 1.145 | 1.055 | 1.386 | 1.070 |
| | Position × Team | 1.124 | 1.041 | 1.375 | 1.060 |
| LIMB-SMN | Team | 1.069 | 1.011 | 1.421 | 1.034 |
| | Position | 1.064 | 1.009 | 1.445 | 1.031 |
| | Sets | 1.102 | 1.028 | 1.374 | 1.050 |
| | Scores | 1.146 | 1.055 | 1.386 | 1.071 |
| | Position × Team | 1.079 | 1.016 | 1.396 | 1.039 |
| LIMB-VAN | Team | 1.176 | 1.076 | 1.410 | 1.085 |
| | Position | 1.060 | 1.008 | 1.470 | 1.029 |
| | Sets | 1.099 | 1.026 | 1.374 | 1.049 |
| | Scores | 1.145 | 1.055 | 1.386 | 1.070 |
| | Position × Team | 1.179 | 1.078 | 1.413 | 1.086 |
| LIMB-VIS | Team | 1.108 | 1.031 | 1.373 | 1.053 |
| | Position | 1.061 | 1.008 | 1.460 | 1.030 |

| | | | | | |
|---|---|---|---|---|---|
| | Sets | 1.104 | 1.029 | 1.373 | 1.050 |
| | Scores | 1.147 | 1.056 | 1.387 | 1.071 |
| | Position × Team | 1.116 | 1.036 | 1.373 | 1.056 |
| SMN-SMN | Team | 1.047 | 1.004 | 1.605 | 1.023 |
| | Position | 1.067 | 1.010 | 1.432 | 1.033 |
| | Sets | 1.097 | 1.025 | 1.376 | 1.047 |
| | Scores | 1.143 | 1.053 | 1.385 | 1.069 |
| | Position × Team | 1.058 | 1.007 | 1.482 | 1.029 |
| SMN-VAN | Team | 1.068 | 1.011 | 1.425 | 1.034 |
| | Position | 1.064 | 1.009 | 1.443 | 1.032 |
| | Sets | 1.100 | 1.027 | 1.374 | 1.049 |
| | Scores | 1.145 | 1.055 | 1.386 | 1.070 |
| | Position × Team | 1.078 | 1.015 | 1.398 | 1.038 |
| SMN-VIS | Team | 1.125 | 1.042 | 1.376 | 1.061 |
| | Position | 1.061 | 1.008 | 1.461 | 1.030 |
| | Sets | 1.101 | 1.027 | 1.374 | 1.049 |
| | Scores | 1.146 | 1.055 | 1.386 | 1.070 |
| | Position × Team | 1.132 | 1.046 | 1.378 | 1.064 |
| VAN-VAN | Team | 1.114 | 1.035 | 1.373 | 1.056 |
| | Position | 1.060 | 1.008 | 1.468 | 1.030 |
| | Sets | 1.107 | 1.031 | 1.373 | 1.052 |
| | Scores | 1.149 | 1.057 | 1.389 | 1.072 |
| | Position × Team | 1.121 | 1.039 | 1.374 | 1.059 |

| | | | | | |
|---|---|---|---|---|---|
| VAN-VIS | Team | 1.112 | 1.034 | 1.373 | 1.055 |
| | Position | 1.061 | 1.008 | 1.463 | 1.030 |
| | Sets | 1.104 | 1.029 | 1.373 | 1.051 |
| | Scores | 1.148 | 1.056 | 1.387 | 1.071 |
| | Position × Team | 1.120 | 1.038 | 1.374 | 1.058 |
| VIS-VIS | Team | 1.061 | 1.008 | 1.463 | 1.030 |
| | Position | 1.065 | 1.010 | 1.440 | 1.032 |
| | Sets | 1.100 | 1.027 | 1.374 | 1.049 |
| | Scores | 1.145 | 1.054 | 1.386 | 1.070 |
| | Position × Team | 1.071 | 1.012 | 1.417 | 1.035 |

**Table S2-1** Full linear mixed-effects model results for associations between network-level functional connectivity differences and Position, Team, and Position × Team in the resting-state condition.

| Brain network(s) | Predictor | *β* | SE | P | P (FDR-corrected) |
|---|---|---|---|---|---|
| DAN-DAN | Position | -0.001 | 0.015 | 0.946 | 0.969 |
| | Team | 0.006 | 0.017 | 0.612 | 0.830 |
| | Position × Team | -0.003 | 0.009 | 0.605 | 0.830 |
| DAN-DMN | Position | -0.004 | 0.010 | 0.536 | 0.804 |
| | Team | -0.005 | 0.011 | 0.521 | 0.796 |
| | Position × Team | 0.001 | 0.006 | 0.748 | 0.898 |
| DAN-FPN | Position | -0.003 | 0.014 | 0.797 | 0.903 |
| | Team | 0.002 | 0.016 | 0.859 | 0.925 |
| | Position × Team | -0.001 | 0.009 | 0.817 | 0.903 |
| DAN-LIMB | Position | -0.009 | 0.013 | 0.325 | 0.690 |
| | Team | -0.014 | 0.015 | 0.204 | 0.600 |
| | Position × Team | 0.007 | 0.008 | 0.256 | 0.614 |
| DAN-SMN | Position | 0.012 | 0.010 | 0.107 | 0.600 |
| | Team | 0.012 | 0.012 | 0.144 | 0.600 |
| | Position × Team | -0.007 | 0.006 | 0.119 | 0.600 |
| DAN-VAN | Position | 0.003 | 0.013 | 0.771 | 0.903 |
| | Team | 0.002 | 0.015 | 0.852 | 0.925 |
| | Position × Team | -0.002 | 0.008 | 0.671 | 0.854 |
| DAN-VIS | Position | -0.010 | 0.010 | 0.145 | 0.600 |
| | Team | -0.004 | 0.011 | 0.585 | 0.830 |

| | | | | | |
|---|---|---|---|---|---|
| | Position × Team | 0.005 | 0.006 | 0.243 | 0.614 |
| DMN-DMN | Position | -0.002 | 0.010 | 0.817 | 0.903 |
| | Team | 0.002 | 0.011 | 0.777 | 0.903 |
| | Position × Team | 0.001 | 0.006 | 0.815 | 0.903 |
| DMN-FPN | Position | -0.007 | 0.011 | 0.381 | 0.696 |
| | Team | 0.001 | 0.012 | 0.886 | 0.936 |
| | Position × Team | 0.002 | 0.007 | 0.642 | 0.843 |
| DMN-LIMB | Position | -0.017 | 0.014 | 0.096 | 0.600 |
| | Team | -0.011 | 0.016 | 0.330 | 0.690 |
| | Position × Team | 0.010 | 0.009 | 0.110 | 0.600 |
| DMN-SMN | Position | -0.006 | 0.009 | 0.367 | 0.692 |
| | Team | -0.004 | 0.011 | 0.594 | 0.830 |
| | Position × Team | 0.002 | 0.006 | 0.586 | 0.830 |
| DMN-VAN | Position | -0.005 | 0.011 | 0.509 | 0.791 |
| | Team | -0.004 | 0.012 | 0.662 | 0.854 |
| | Position × Team | 0.002 | 0.007 | 0.632 | 0.843 |
| DMN-VIS | Position | -0.011 | 0.010 | 0.114 | 0.600 |
| | Team | -0.003 | 0.011 | 0.732 | 0.892 |
| | Position × Team | 0.005 | 0.006 | 0.221 | 0.600 |
| FPN-FPN | Position | 0.032 | 0.017 | 0.009 | 0.130 |
| | Team | 0.041 | 0.019 | 0.003 | 0.088 |
| | Position × Team | -0.021 | 0.011 | 0.004 | 0.088 |
| FPN-LIMB | Position | 0.009 | 0.013 | 0.336 | 0.690 |

| | | | | | |
|---|---|---|---|---|---|
| | Team | 0.012 | 0.014 | 0.217 | 0.600 |
| | Position × Team | -0.004 | 0.008 | 0.437 | 0.735 |
| FPN-SMN | Position | 0.009 | 0.010 | 0.195 | 0.600 |
| | Team | 0.011 | 0.011 | 0.163 | 0.600 |
| | Position × Team | -0.007 | 0.006 | 0.112 | 0.600 |
| FPN-VAN | Position | 0.008 | 0.012 | 0.358 | 0.692 |
| | Team | 0.017 | 0.014 | 0.087 | 0.600 |
| | Position × Team | -0.006 | 0.008 | 0.228 | 0.600 |
| FPN-VIS | Position | -0.007 | 0.011 | 0.393 | 0.704 |
| | Team | -0.001 | 0.013 | 0.894 | 0.936 |
| | Position × Team | 0.003 | 0.007 | 0.500 | 0.791 |
| LIMB-LIMB | Position | 0.018 | 0.025 | 0.310 | 0.689 |
| | Team | 0.018 | 0.028 | 0.371 | 0.692 |
| | Position × Team | -0.011 | 0.015 | 0.292 | 0.682 |
| LIMB-SMN | Position | -0.017 | 0.016 | 0.144 | 0.600 |
| | Team | -0.016 | 0.019 | 0.217 | 0.600 |
| | Position × Team | 0.008 | 0.010 | 0.253 | 0.614 |
| LIMB-VAN | Position | -0.016 | 0.018 | 0.196 | 0.600 |
| | Team | -0.013 | 0.020 | 0.360 | 0.692 |
| | Position × Team | 0.009 | 0.011 | 0.222 | 0.600 |
| LIMB-VIS | Position | -0.011 | 0.011 | 0.163 | 0.600 |
| | Team | -0.012 | 0.013 | 0.192 | 0.600 |
| | Position × Team | 0.006 | 0.007 | 0.208 | 0.600 |

| | | | | | |
|---|---|---|---|---|---|
| SMN-SMN | Position | 0.040 | 0.015 | < 0.001 | **0.012** |
| | Team | 0.034 | 0.017 | 0.004 | 0.088 |
| | Position × Team | -0.017 | 0.009 | 0.009 | 0.130 |
| SMN-VAN | Position | -0.018 | 0.012 | 0.040 | 0.422 |
| | Team | -0.020 | 0.014 | 0.040 | 0.422 |
| | Position × Team | 0.010 | 0.007 | 0.064 | 0.600 |
| SMN-VIS | Position | -0.006 | 0.011 | 0.431 | 0.735 |
| | Team | < 0.001 | 0.012 | 0.980 | 0.980 |
| | Position × Team | 0.002 | 0.007 | 0.714 | 0.882 |
| VAN-VAN | Position | -0.002 | 0.019 | 0.902 | 0.936 |
| | Team | -0.008 | 0.021 | 0.585 | 0.830 |
| | Position × Team | 0.003 | 0.012 | 0.696 | 0.872 |
| VAN-VIS | Position | -0.012 | 0.011 | 0.143 | 0.600 |
| | Team | -0.007 | 0.013 | 0.435 | 0.735 |
| | Position × Team | 0.005 | 0.007 | 0.311 | 0.689 |
| VIS-VIS | Position | < 0.001 | 0.011 | 0.961 | 0.972 |
| | Team | 0.006 | 0.013 | 0.484 | 0.791 |
| | Position × Team | -0.003 | 0.007 | 0.504 | 0.791 |

**Table S2-2** Full linear mixed-effects model results for associations between network-level functional connectivity differences and Position, Team, and Position × Team in the resting-state condition, controlling for Sets and Scores.

| Brain network(s) | Predictor | $\beta$ | SE | P | P (FDR-corrected) |
|---|---|---|---|---|---|
| DAN-DAN | Position | -0.001 | 0.015 | 0.957 | 0.988 |
| | Team | 0.006 | 0.017 | 0.603 | 0.898 |
| | Position × Team | -0.001 | 0.003 | 0.526 | 0.883 |
| | Sets | < 0.001 | 0.003 | 0.902 | 0.988 |
| | Scores | -0.004 | 0.009 | 0.596 | 0.898 |
| DAN-DMN | Position | -0.005 | 0.010 | 0.500 | 0.876 |
| | Team | -0.005 | 0.011 | 0.502 | 0.876 |
| | Position × Team | 0.001 | 0.002 | 0.280 | 0.741 |
| | Sets | 0.001 | 0.002 | 0.543 | 0.885 |
| | Scores | 0.001 | 0.006 | 0.724 | 0.933 |
| DAN-FPN | Position | -0.002 | 0.014 | 0.820 | 0.973 |
| | Team | 0.002 | 0.016 | 0.855 | 0.973 |
| | Position × Team | < 0.001 | 0.002 | 0.935 | 0.988 |
| | Sets | -0.001 | 0.003 | 0.720 | 0.933 |
| | Scores | -0.001 | 0.009 | 0.812 | 0.971 |
| DAN-LIMB | Position | -0.009 | 0.013 | 0.365 | 0.808 |
| | Team | -0.014 | 0.015 | 0.211 | 0.732 |
| | Position × Team | -0.001 | 0.002 | 0.719 | 0.933 |
| | Sets | -0.002 | 0.003 | 0.375 | 0.808 |
| | Scores | 0.007 | 0.008 | 0.266 | 0.741 |

| | | | | | |
|---|---|---|---|---|---|
| DAN-SMN | Position | 0.012 | 0.011 | 0.108 | 0.732 |
| | Team | 0.012 | 0.012 | 0.140 | 0.732 |
| | Position × Team | -0.001 | 0.002 | 0.426 | 0.841 |
| | Sets | < 0.001 | 0.002 | 0.992 | 0.992 |
| | Scores | -0.007 | 0.006 | 0.115 | 0.732 |
| DAN-VAN | Position | 0.003 | 0.013 | 0.775 | 0.964 |
| | Team | 0.002 | 0.015 | 0.847 | 0.973 |
| | Position × Team | -0.001 | 0.002 | 0.679 | 0.924 |
| | Sets | < 0.001 | 0.003 | 0.941 | 0.988 |
| | Scores | -0.002 | 0.008 | 0.667 | 0.924 |
| DAN-VIS | Position | -0.010 | 0.010 | 0.150 | 0.732 |
| | Team | -0.004 | 0.011 | 0.594 | 0.898 |
| | Position × Team | -0.001 | 0.002 | 0.602 | 0.898 |
| | Sets | < 0.001 | 0.002 | 0.931 | 0.988 |
| | Scores | 0.005 | 0.006 | 0.249 | 0.741 |
| DMN-DMN | Position | -0.002 | 0.010 | 0.798 | 0.964 |
| | Team | 0.002 | 0.011 | 0.799 | 0.964 |
| | Position × Team | 0.001 | 0.002 | 0.235 | 0.732 |
| | Sets | < 0.001 | 0.002 | 0.834 | 0.973 |
| | Scores | 0.001 | 0.006 | 0.793 | 0.964 |
| DMN-FPN | Position | -0.007 | 0.011 | 0.350 | 0.804 |
| | Team | 0.001 | 0.012 | 0.897 | 0.988 |
| | Position × Team | < 0.001 | 0.002 | 0.791 | 0.964 |

| | | | | | |
|---|---|---|---|---|---|
| | Sets | 0.001 | 0.002 | 0.457 | 0.866 |
| | Scores | 0.002 | 0.007 | 0.631 | 0.902 |
| DMN-LIMB | Position | -0.017 | 0.014 | 0.094 | 0.732 |
| | Team | -0.011 | 0.016 | 0.325 | 0.804 |
| | Position × Team | 0.001 | 0.002 | 0.625 | 0.902 |
| | Sets | < 0.001 | 0.003 | 0.884 | 0.988 |
| | Scores | 0.010 | 0.009 | 0.108 | 0.732 |
| DMN-SMN | Position | -0.006 | 0.009 | 0.342 | 0.804 |
| | Team | -0.004 | 0.011 | 0.599 | 0.898 |
| | Position × Team | -0.001 | 0.002 | 0.490 | 0.876 |
| | Sets | 0.001 | 0.002 | 0.492 | 0.876 |
| | Scores | 0.002 | 0.006 | 0.589 | 0.898 |
| DMN-VAN | Position | -0.005 | 0.011 | 0.507 | 0.876 |
| | Team | -0.004 | 0.012 | 0.671 | 0.924 |
| | Position × Team | -0.001 | 0.002 | 0.536 | 0.883 |
| | Sets | < 0.001 | 0.002 | 0.902 | 0.988 |
| | Scores | 0.002 | 0.007 | 0.641 | 0.906 |
| DMN-VIS | Position | -0.011 | 0.010 | 0.099 | 0.732 |
| | Team | -0.003 | 0.011 | 0.733 | 0.933 |
| | Position × Team | -0.001 | 0.002 | 0.564 | 0.898 |
| | Sets | 0.002 | 0.002 | 0.329 | 0.804 |
| | Scores | 0.005 | 0.006 | 0.220 | 0.732 |
| FPN-FPN | Position | 0.032 | 0.017 | 0.009 | 0.223 |

| | | | | | |
|---|---|---|---|---|---|
| | Team | 0.042 | 0.019 | 0.002 | 0.148 |
| | Position × Team | -0.003 | 0.003 | 0.150 | 0.732 |
| | Sets | < 0.001 | 0.004 | 0.904 | 0.988 |
| | Scores | -0.022 | 0.011 | 0.004 | 0.148 |
| FPN-LIMB | Position | 0.008 | 0.013 | 0.350 | 0.804 |
| | Team | 0.013 | 0.014 | 0.207 | 0.732 |
| | Position × Team | -0.003 | 0.002 | 0.096 | 0.732 |
| | Sets | 0.001 | 0.003 | 0.675 | 0.924 |
| | Scores | -0.004 | 0.008 | 0.423 | 0.841 |
| FPN-SMN | Position | 0.010 | 0.010 | 0.163 | 0.732 |
| | Team | 0.012 | 0.011 | 0.152 | 0.732 |
| | Position × Team | -0.001 | 0.002 | 0.276 | 0.741 |
| | Sets | -0.002 | 0.002 | 0.244 | 0.741 |
| | Scores | -0.007 | 0.006 | 0.102 | 0.732 |
| FPN-VAN | Position | 0.007 | 0.012 | 0.390 | 0.828 |
| | Team | 0.017 | 0.014 | 0.083 | 0.732 |
| | Position × Team | -0.002 | 0.002 | 0.132 | 0.732 |
| | Sets | 0.002 | 0.003 | 0.417 | 0.841 |
| | Scores | -0.007 | 0.008 | 0.222 | 0.732 |
| FPN-VIS | Position | -0.006 | 0.011 | 0.423 | 0.841 |
| | Team | -0.001 | 0.013 | 0.922 | 0.988 |
| | Position × Team | -0.002 | 0.002 | 0.158 | 0.732 |
| | Sets | -0.001 | 0.003 | 0.615 | 0.898 |

| | | | | | |
|---|---|---|---|---|---|
| | Scores | 0.003 | 0.007 | 0.523 | 0.883 |
| LIMB-LIMB | Position | 0.018 | 0.025 | 0.305 | 0.792 |
| | Team | 0.018 | 0.028 | 0.356 | 0.804 |
| | Position × Team | -0.004 | 0.004 | 0.155 | 0.732 |
| | Sets | < 0.001 | 0.005 | 0.950 | 0.988 |
| | Scores | -0.012 | 0.015 | 0.280 | 0.741 |
| LIMB-SMN | Position | -0.017 | 0.017 | 0.149 | 0.732 |
| | Team | -0.016 | 0.019 | 0.228 | 0.732 |
| | Position × Team | -0.002 | 0.003 | 0.225 | 0.732 |
| | Sets | < 0.001 | 0.004 | 0.987 | 0.992 |
| | Scores | 0.008 | 0.010 | 0.264 | 0.741 |
| LIMB-VAN | Position | -0.016 | 0.018 | 0.210 | 0.732 |
| | Team | -0.013 | 0.020 | 0.371 | 0.808 |
| | Position × Team | -0.002 | 0.003 | 0.443 | 0.852 |
| | Sets | -0.001 | 0.004 | 0.721 | 0.933 |
| | Scores | 0.009 | 0.011 | 0.231 | 0.732 |
| LIMB-VIS | Position | -0.010 | 0.011 | 0.196 | 0.732 |
| | Team | -0.012 | 0.013 | 0.189 | 0.732 |
| | Position × Team | 0.001 | 0.002 | 0.278 | 0.741 |
| | Sets | -0.002 | 0.003 | 0.195 | 0.732 |
| | Scores | 0.006 | 0.007 | 0.207 | 0.732 |
| SMN-SMN | Position | 0.039 | 0.015 | < 0.001 | **0.032** |
| | Team | 0.034 | 0.017 | 0.004 | 0.148 |

| | | | | | |
|---|---|---|---|---|---|
| | Position × Team | -0.001 | 0.003 | 0.536 | 0.883 |
| | Sets | 0.003 | 0.003 | 0.144 | 0.732 |
| | Scores | -0.017 | 0.009 | 0.009 | 0.223 |
| SMN-VAN | Position | -0.016 | 0.012 | 0.054 | 0.732 |
| | Team | -0.020 | 0.014 | 0.036 | 0.633 |
| | Position × Team | 0.004 | 0.002 | 0.012 | 0.245 |
| | Sets | -0.004 | 0.003 | 0.055 | 0.732 |
| | Scores | 0.010 | 0.007 | 0.060 | 0.732 |
| SMN-VIS | Position | -0.006 | 0.011 | 0.420 | 0.841 |
| | Team | < 0.001 | 0.012 | 0.963 | 0.988 |
| | Position × Team | 0.001 | 0.002 | 0.355 | 0.804 |
| | Sets | < 0.001 | 0.003 | 0.849 | 0.973 |
| | Scores | 0.002 | 0.007 | 0.698 | 0.933 |
| VAN-VAN | Position | < 0.001 | 0.019 | 0.990 | 0.992 |
| | Team | -0.008 | 0.021 | 0.613 | 0.898 |
| | Position × Team | -0.003 | 0.003 | 0.227 | 0.732 |
| | Sets | -0.004 | 0.004 | 0.212 | 0.732 |
| | Scores | 0.003 | 0.012 | 0.730 | 0.933 |
| VAN-VIS | Position | -0.011 | 0.011 | 0.153 | 0.732 |
| | Team | -0.007 | 0.013 | 0.444 | 0.852 |
| | Position × Team | -0.001 | 0.002 | 0.603 | 0.898 |
| | Sets | -0.001 | 0.003 | 0.753 | 0.950 |
| | Scores | 0.005 | 0.007 | 0.319 | 0.804 |

| | | | | | |
|---|---|---|---|---|---|
| VIS-VIS | Position | < 0.001 | 0.011 | 0.967 | 0.988 |
| | Team | 0.006 | 0.013 | 0.482 | 0.876 |
| | Position × Team | < 0.001 | 0.002 | 0.837 | 0.973 |
| | Sets | < 0.001 | 0.003 | 0.943 | 0.988 |
| | Scores | -0.003 | 0.007 | 0.501 | 0.876 |

**Table S2-3** Full linear mixed-effects model results for associations between network-level functional connectivity differences and Position, Team, and Position × Team in the game-viewing condition.

| Brain network(s) | Predictor | *β* | SE | P | P (FDR-corrected) |
|---|---|---|---|---|---|
| DAN-DAN | Position | -0.005 | 0.012 | 0.508 | 0.562 |
| | Team | -0.005 | 0.013 | 0.616 | 0.655 |
| | Position × Team | 0.001 | 0.007 | 0.823 | 0.843 |
| DAN-DMN | Position | 0.007 | 0.009 | 0.262 | 0.319 |
| | Team | 0.010 | 0.010 | 0.139 | 0.186 |
| | Position × Team | -0.006 | 0.005 | 0.101 | 0.144 |
| DAN-FPN | Position | -0.004 | 0.010 | 0.586 | 0.632 |
| | Team | 0.004 | 0.011 | 0.580 | 0.632 |
| | Position × Team | < 0.001 | 0.006 | 0.918 | 0.918 |
| DAN-LIMB | Position | 0.013 | 0.012 | 0.125 | 0.170 |
| | Team | 0.009 | 0.013 | 0.350 | 0.420 |
| | Position × Team | -0.006 | 0.007 | 0.206 | 0.263 |
| DAN-SMN | Position | 0.020 | 0.008 | 0.001 | **0.002** |
| | Team | 0.020 | 0.009 | 0.002 | **0.007** |
| | Position × Team | -0.012 | 0.005 | 0.001 | **0.004** |
| DAN-VAN | Position | 0.016 | 0.010 | 0.023 | **0.040** |
| | Team | 0.021 | 0.011 | 0.012 | **0.023** |
| | Position × Team | -0.012 | 0.006 | 0.006 | **0.014** |
| DAN-VIS | Position | 0.018 | 0.012 | 0.037 | 0.058 |
| | Team | 0.015 | 0.014 | 0.125 | 0.170 |

| | | | | | |
|---|---|---|---|---|---|
| | Position × Team | -0.010 | 0.007 | 0.060 | 0.093 |
| DMN-DMN | Position | 0.007 | 0.012 | 0.379 | 0.443 |
| | Team | 0.009 | 0.013 | 0.362 | 0.428 |
| | Position × Team | -0.006 | 0.007 | 0.236 | 0.296 |
| DMN-FPN | Position | 0.022 | 0.015 | 0.032 | 0.052 |
| | Team | 0.030 | 0.017 | 0.012 | **0.023** |
| | Position × Team | -0.018 | 0.009 | 0.005 | **0.012** |
| DMN-LIMB | Position | 0.046 | 0.015 | < 0.001 | **< 0.001** |
| | Team | 0.054 | 0.017 | < 0.001 | **< 0.001** |
| | Position × Team | -0.029 | 0.009 | < 0.001 | **< 0.001** |
| DMN-SMN | Position | 0.025 | 0.010 | < 0.001 | **0.002** |
| | Team | 0.033 | 0.011 | < 0.001 | **< 0.001** |
| | Position × Team | -0.019 | 0.006 | < 0.001 | **< 0.001** |
| DMN-VAN | Position | 0.034 | 0.014 | < 0.001 | **0.002** |
| | Team | 0.034 | 0.016 | 0.002 | **0.007** |
| | Position × Team | -0.020 | 0.008 | 0.001 | **0.004** |
| DMN-VIS | Position | 0.036 | 0.010 | < 0.001 | **< 0.001** |
| | Team | 0.043 | 0.011 | < 0.001 | **< 0.001** |
| | Position × Team | -0.025 | 0.006 | < 0.001 | **< 0.001** |
| FPN-FPN | Position | 0.031 | 0.019 | 0.020 | **0.036** |
| | Team | 0.038 | 0.021 | 0.012 | **0.023** |
| | Position × Team | -0.023 | 0.012 | 0.004 | **0.011** |
| FPN-LIMB | Position | 0.028 | 0.014 | 0.005 | **0.012** |

| | | | | | |
|---|---|---|---|---|---|
| | Team | 0.021 | 0.016 | 0.065 | 0.096 |
| | Position × Team | -0.013 | 0.009 | 0.032 | 0.052 |
| FPN-SMN | Position | 0.019 | 0.011 | 0.015 | **0.029** |
| | Team | 0.025 | 0.013 | 0.005 | **0.012** |
| | Position × Team | -0.015 | 0.007 | 0.002 | **0.005** |
| FPN-VAN | Position | 0.031 | 0.016 | 0.005 | **0.012** |
| | Team | 0.030 | 0.018 | 0.019 | **0.036** |
| | Position × Team | -0.019 | 0.010 | 0.007 | **0.014** |
| FPN-VIS | Position | 0.045 | 0.011 | < 0.001 | **< 0.001** |
| | Team | 0.051 | 0.012 | < 0.001 | **< 0.001** |
| | Position × Team | -0.030 | 0.007 | < 0.001 | **< 0.001** |
| LIMB-LIMB | Position | 0.015 | 0.019 | 0.248 | 0.307 |
| | Team | 0.026 | 0.022 | 0.094 | 0.137 |
| | Position × Team | -0.013 | 0.012 | 0.118 | 0.166 |
| LIMB-SMN | Position | 0.017 | 0.013 | 0.062 | 0.094 |
| | Team | 0.022 | 0.015 | 0.031 | 0.052 |
| | Position × Team | -0.012 | 0.008 | 0.026 | **0.045** |
| LIMB-VAN | Position | 0.034 | 0.015 | 0.002 | **0.007** |
| | Team | 0.034 | 0.018 | 0.007 | **0.014** |
| | Position × Team | -0.020 | 0.010 | 0.003 | **0.008** |
| LIMB-VIS | Position | 0.005 | 0.011 | 0.505 | 0.562 |
| | Team | 0.002 | 0.013 | 0.833 | 0.843 |
| | Position × Team | -0.002 | 0.007 | 0.732 | 0.760 |

| | | | | | |
|---|---|---|---|---|---|
| SMN-SMN | Position | 0.037 | 0.011 | < 0.001 | **< 0.001** |
| | Team | 0.034 | 0.012 | < 0.001 | **0.001** |
| | Position × Team | -0.019 | 0.007 | < 0.001 | **< 0.001** |
| SMN-VAN | Position | 0.030 | 0.012 | < 0.001 | **0.002** |
| | Team | 0.036 | 0.014 | < 0.001 | **0.001** |
| | Position × Team | -0.019 | 0.007 | < 0.001 | **0.002** |
| SMN-VIS | Position | 0.037 | 0.010 | < 0.001 | **< 0.001** |
| | Team | 0.041 | 0.012 | < 0.001 | **< 0.001** |
| | Position × Team | -0.024 | 0.006 | < 0.001 | **< 0.001** |
| VAN-VAN | Position | 0.029 | 0.019 | 0.034 | 0.055 |
| | Team | 0.022 | 0.022 | 0.156 | 0.205 |
| | Position × Team | -0.011 | 0.012 | 0.177 | 0.230 |
| VAN-VIS | Position | 0.005 | 0.011 | 0.469 | 0.540 |
| | Team | 0.003 | 0.012 | 0.725 | 0.760 |
| | Position × Team | -0.003 | 0.007 | 0.498 | 0.562 |
| VIS-VIS | Position | 0.030 | 0.012 | 0.001 | **0.002** |
| | Team | 0.030 | 0.014 | 0.003 | **0.007** |
| | Position × Team | -0.018 | 0.008 | 0.001 | **0.003** |

**Table S2-4** Full linear mixed-effects model results for associations between network-level functional connectivity differences and Position, Team, and Position × Team in the game-viewing condition, controlling for Sets and Scores.

| Brain network(s) | Predictor | *β* | SE | P | P (FDR-corrected) |
|---|---|---|---|---|---|
| DAN-DAN | Position | -0.004 | 0.011 | 0.600 | 0.757 |
| | Team | -0.005 | 0.013 | 0.595 | 0.757 |
| | Position × Team | 0.004 | 0.002 | 0.010 | **0.033** |
| | Sets | -0.003 | 0.003 | 0.058 | 0.134 |
| | Scores | 0.001 | 0.007 | 0.810 | 0.900 |
| DAN-DMN | Position | 0.007 | 0.009 | 0.217 | 0.358 |
| | Team | 0.010 | 0.010 | 0.143 | 0.260 |
| | Position × Team | 0.002 | 0.001 | 0.076 | 0.166 |
| | Sets | -0.002 | 0.002 | 0.130 | 0.244 |
| | Scores | -0.006 | 0.005 | 0.102 | 0.214 |
| DAN-FPN | Position | -0.003 | 0.010 | 0.634 | 0.776 |
| | Team | 0.004 | 0.011 | 0.581 | 0.753 |
| | Position × Team | 0.001 | 0.002 | 0.597 | 0.757 |
| | Sets | -0.001 | 0.002 | 0.401 | 0.579 |
| | Scores | < 0.001 | 0.006 | 0.915 | 0.949 |
| DAN-LIMB | Position | 0.013 | 0.012 | 0.122 | 0.241 |
| | Team | 0.009 | 0.013 | 0.351 | 0.529 |
| | Position × Team | < 0.001 | 0.002 | 0.845 | 0.913 |
| | Sets | -0.001 | 0.003 | 0.776 | 0.888 |
| | Scores | -0.006 | 0.007 | 0.206 | 0.345 |

| | | | | | |
|---|---|---|---|---|---|
| DAN-SMN | Position | 0.020 | 0.008 | 0.001 | **0.004** |
| | Team | 0.020 | 0.009 | 0.002 | **0.011** |
| | Position × Team | < 0.001 | 0.001 | 0.664 | 0.795 |
| | Sets | < 0.001 | 0.002 | 0.938 | 0.966 |
| | Scores | -0.012 | 0.005 | 0.001 | **0.006** |
| DAN-VAN | Position | 0.017 | 0.010 | 0.020 | 0.055 |
| | Team | 0.021 | 0.011 | 0.012 | **0.037** |
| | Position × Team | 0.001 | 0.002 | 0.478 | 0.657 |
| | Sets | -0.001 | 0.002 | 0.465 | 0.645 |
| | Scores | -0.012 | 0.006 | 0.006 | **0.022** |
| DAN-VIS | Position | 0.016 | 0.012 | 0.055 | 0.129 |
| | Team | 0.014 | 0.014 | 0.132 | 0.244 |
| | Position × Team | < 0.001 | 0.002 | 0.958 | 0.972 |
| | Sets | 0.004 | 0.003 | 0.050 | 0.120 |
| | Scores | -0.010 | 0.007 | 0.065 | 0.146 |
| DMN-DMN | Position | 0.007 | 0.012 | 0.390 | 0.575 |
| | Team | 0.009 | 0.013 | 0.361 | 0.539 |
| | Position × Team | < 0.001 | 0.002 | 0.782 | 0.888 |
| | Sets | < 0.001 | 0.003 | 0.839 | 0.913 |
| | Scores | -0.006 | 0.007 | 0.236 | 0.385 |
| DMN-FPN | Position | 0.022 | 0.015 | 0.035 | 0.086 |
| | Team | 0.030 | 0.017 | 0.011 | **0.036** |
| | Position × Team | -0.003 | 0.002 | 0.125 | 0.241 |

| | | | | | |
|---|---|---|---|---|---|
| | Sets | 0.001 | 0.003 | 0.644 | 0.778 |
| | Scores | -0.018 | 0.009 | 0.005 | **0.018** |
| DMN-LIMB | Position | 0.047 | 0.015 | < 0.001 | **< 0.001** |
| | Team | 0.054 | 0.017 | < 0.001 | **< 0.001** |
| | Position × Team | -0.001 | 0.002 | 0.446 | 0.629 |
| | Sets | -0.002 | 0.003 | 0.281 | 0.437 |
| | Scores | -0.029 | 0.009 | < 0.001 | **< 0.001** |
| DMN-SMN | Position | 0.024 | 0.010 | 0.001 | **0.004** |
| | Team | 0.033 | 0.011 | < 0.001 | **< 0.001** |
| | Position × Team | -0.003 | 0.002 | 0.015 | **0.046** |
| | Sets | 0.002 | 0.002 | 0.124 | 0.241 |
| | Scores | -0.019 | 0.006 | < 0.001 | **< 0.001** |
| DMN-VAN | Position | 0.033 | 0.014 | 0.001 | **0.004** |
| | Team | 0.034 | 0.016 | 0.002 | **0.011** |
| | Position × Team | -0.002 | 0.002 | 0.205 | 0.345 |
| | Sets | 0.002 | 0.003 | 0.265 | 0.423 |
| | Scores | -0.020 | 0.009 | 0.001 | **0.006** |
| DMN-VIS | Position | 0.036 | 0.010 | < 0.001 | **< 0.001** |
| | Team | 0.043 | 0.011 | < 0.001 | **< 0.001** |
| | Position × Team | -0.001 | 0.002 | 0.275 | 0.434 |
| | Sets | < 0.001 | 0.002 | 0.848 | 0.913 |
| | Scores | -0.025 | 0.006 | < 0.001 | **< 0.001** |
| FPN-FPN | Position | 0.028 | 0.019 | 0.038 | 0.092 |

| | | | | | |
|---|---|---|---|---|---|
| | Team | 0.037 | 0.021 | 0.014 | **0.043** |
| | Position × Team | -0.001 | 0.003 | 0.618 | 0.773 |
| | Sets | 0.007 | 0.004 | 0.006 | **0.022** |
| | Scores | -0.023 | 0.012 | 0.005 | **0.021** |
| FPN-LIMB | Position | 0.027 | 0.014 | 0.007 | **0.025** |
| | Team | 0.021 | 0.016 | 0.069 | 0.151 |
| | Position × Team | < 0.001 | 0.002 | 0.878 | 0.925 |
| | Sets | 0.003 | 0.003 | 0.183 | 0.317 |
| | Scores | -0.013 | 0.009 | 0.035 | 0.086 |
| FPN-SMN | Position | 0.019 | 0.011 | 0.016 | **0.046** |
| | Team | 0.025 | 0.013 | 0.005 | **0.019** |
| | Position × Team | -0.001 | 0.002 | 0.303 | 0.467 |
| | Sets | < 0.001 | 0.003 | 0.887 | 0.926 |
| | Scores | -0.016 | 0.007 | 0.001 | **0.008** |
| FPN-VAN | Position | 0.032 | 0.016 | 0.005 | **0.018** |
| | Team | 0.030 | 0.018 | 0.019 | 0.052 |
| | Position × Team | -0.001 | 0.003 | 0.636 | 0.776 |
| | Sets | -0.001 | 0.003 | 0.578 | 0.753 |
| | Scores | -0.019 | 0.010 | 0.007 | **0.022** |
| FPN-VIS | Position | 0.045 | 0.011 | < 0.001 | **< 0.001** |
| | Team | 0.051 | 0.012 | < 0.001 | **< 0.001** |
| | Position × Team | -0.002 | 0.002 | 0.122 | 0.241 |
| | Sets | < 0.001 | 0.002 | 0.997 | 0.997 |

| | | | | | |
|---|---|---|---|---|---|
| | Scores | -0.030 | 0.007 | < 0.001 | **< 0.001** |
| LIMB-LIMB | Position | 0.015 | 0.019 | 0.256 | 0.413 |
| | Team | 0.025 | 0.022 | 0.096 | 0.204 |
| | Position × Team | < 0.001 | 0.003 | 0.955 | 0.972 |
| | Sets | < 0.001 | 0.004 | 0.867 | 0.919 |
| | Scores | -0.013 | 0.012 | 0.119 | 0.241 |
| LIMB-SMN | Position | 0.017 | 0.013 | 0.059 | 0.134 |
| | Team | 0.023 | 0.015 | 0.028 | 0.072 |
| | Position × Team | -0.002 | 0.002 | 0.130 | 0.244 |
| | Sets | -0.001 | 0.003 | 0.800 | 0.896 |
| | Scores | -0.013 | 0.008 | 0.024 | 0.064 |
| LIMB-VAN | Position | 0.034 | 0.016 | 0.002 | **0.011** |
| | Team | 0.034 | 0.018 | 0.006 | **0.022** |
| | Position × Team | -0.002 | 0.003 | 0.405 | 0.579 |
| | Sets | < 0.001 | 0.003 | 0.988 | 0.995 |
| | Scores | -0.020 | 0.010 | 0.003 | **0.012** |
| LIMB-VIS | Position | 0.006 | 0.011 | 0.449 | 0.629 |
| | Team | 0.002 | 0.013 | 0.855 | 0.914 |
| | Position × Team | 0.003 | 0.002 | 0.017 | **0.048** |
| | Sets | -0.002 | 0.002 | 0.195 | 0.333 |
| | Scores | -0.002 | 0.007 | 0.747 | 0.879 |
| SMN-SMN | Position | 0.038 | 0.011 | < 0.001 | **< 0.001** |
| | Team | 0.033 | 0.012 | < 0.001 | **0.001** |

| | | | | | |
|---|---|---|---|---|---|
| | Position × Team | 0.004 | 0.002 | 0.003 | **0.012** |
| | Sets | -0.002 | 0.003 | 0.175 | 0.307 |
| | Scores | -0.019 | 0.007 | < 0.001 | **< 0.001** |
| SMN-VAN | Position | 0.030 | 0.012 | < 0.001 | **0.003** |
| | Team | 0.036 | 0.014 | < 0.001 | **0.002** |
| | Position × Team | -0.001 | 0.002 | 0.323 | 0.492 |
| | Sets | 0.001 | 0.003 | 0.787 | 0.888 |
| | Scores | -0.019 | 0.007 | < 0.001 | **0.003** |
| SMN-VIS | Position | 0.037 | 0.010 | < 0.001 | **< 0.001** |
| | Team | 0.041 | 0.012 | < 0.001 | **< 0.001** |
| | Position × Team | -0.001 | 0.002 | 0.526 | 0.701 |
| | Sets | < 0.001 | 0.002 | 0.768 | 0.888 |
| | Scores | -0.024 | 0.006 | < 0.001 | **< 0.001** |
| VAN-VAN | Position | 0.031 | 0.019 | 0.025 | 0.065 |
| | Team | 0.023 | 0.022 | 0.145 | 0.261 |
| | Position × Team | -0.002 | 0.003 | 0.395 | 0.577 |
| | Sets | -0.005 | 0.004 | 0.122 | 0.241 |
| | Scores | -0.012 | 0.012 | 0.163 | 0.290 |
| VAN-VIS | Position | 0.005 | 0.011 | 0.499 | 0.676 |
| | Team | 0.003 | 0.012 | 0.727 | 0.862 |
| | Position × Team | < 0.001 | 0.002 | 0.823 | 0.907 |
| | Sets | 0.001 | 0.002 | 0.569 | 0.752 |
| | Scores | -0.003 | 0.007 | 0.502 | 0.676 |

| | | | | | |
|---|---|---|---|---|---|
| VIS-VIS | Position | 0.031 | 0.012 | 0.001 | **0.004** |
| | Team | 0.030 | 0.014 | 0.003 | **0.012** |
| | Position × Team | < 0.001 | 0.002 | 0.780 | 0.888 |
| | Sets | -0.001 | 0.003 | 0.637 | 0.776 |
| | Scores | -0.018 | 0.008 | 0.001 | **0.005** |

**Table S3-1** Full linear mixed-effects model results for associations between network-level functional connectivity differences and Sets, Team, and Sets × Team in the resting-state condition.

| Brain network(s) | Predictor | *β* | SE | P | P (FDR-corrected) |
|---|---|---|---|---|---|
| DAN-DAN | Sets | 0.001 | 0.006 | 0.828 | 0.940 |
| | Team | < 0.001 | 0.004 | 0.907 | 0.941 |
| | Sets × Team | -0.001 | 0.004 | 0.602 | 0.872 |
| DAN-DMN | Sets | 0.001 | 0.004 | 0.762 | 0.923 |
| | Team | -0.003 | 0.002 | 0.156 | 0.524 |
| | Sets × Team | < 0.001 | 0.002 | 0.781 | 0.923 |
| DAN-FPN | Sets | < 0.001 | 0.006 | 0.907 | 0.941 |
| | Team | < 0.001 | 0.004 | 0.894 | 0.941 |
| | Sets × Team | < 0.001 | 0.003 | 0.848 | 0.941 |
| DAN-LIMB | Sets | -0.008 | 0.005 | 0.045 | 0.201 |
| | Team | -0.002 | 0.003 | 0.451 | 0.790 |
| | Sets × Team | 0.004 | 0.003 | 0.059 | 0.250 |
| DAN-SMN | Sets | -0.004 | 0.004 | 0.227 | 0.672 |
| | Team | -0.001 | 0.003 | 0.773 | 0.923 |
| | Sets × Team | 0.002 | 0.003 | 0.340 | 0.729 |
| DAN-VAN | Sets | 0.002 | 0.005 | 0.534 | 0.847 |
| | Team | -0.002 | 0.003 | 0.327 | 0.724 |
| | Sets × Team | -0.002 | 0.003 | 0.392 | 0.732 |
| DAN-VIS | Sets | -0.002 | 0.004 | 0.586 | 0.872 |
| | Team | 0.005 | 0.002 | 0.008 | 0.088 |

| | | | | | |
|---|---|---|---|---|---|
| | Sets × Team | 0.001 | 0.002 | 0.729 | 0.923 |
| DMN-DMN | Sets | 0.008 | 0.004 | 0.003 | 0.088 |
| | Team | 0.004 | 0.003 | 0.022 | 0.131 |
| | Sets × Team | -0.005 | 0.002 | 0.008 | 0.088 |
| DMN-FPN | Sets | 0.003 | 0.004 | 0.311 | 0.722 |
| | Team | 0.005 | 0.003 | 0.007 | 0.088 |
| | Sets × Team | -0.002 | 0.003 | 0.390 | 0.732 |
| DMN-LIMB | Sets | 0.010 | 0.006 | 0.016 | 0.115 |
| | Team | 0.007 | 0.004 | 0.008 | 0.088 |
| | Sets × Team | -0.006 | 0.004 | 0.017 | 0.115 |
| DMN-SMN | Sets | 0.001 | 0.004 | 0.594 | 0.872 |
| | Team | < 0.001 | 0.002 | 0.949 | 0.972 |
| | Sets × Team | -0.001 | 0.002 | 0.424 | 0.774 |
| DMN-VAN | Sets | 0.001 | 0.004 | 0.862 | 0.941 |
| | Team | < 0.001 | 0.003 | 0.864 | 0.941 |
| | Sets × Team | -0.001 | 0.003 | 0.644 | 0.901 |
| DMN-VIS | Sets | 0.007 | 0.004 | 0.007 | 0.088 |
| | Team | 0.007 | 0.002 | < 0.001 | **0.010** |
| | Sets × Team | -0.005 | 0.002 | 0.002 | 0.088 |
| FPN-FPN | Sets | 0.002 | 0.007 | 0.677 | 0.923 |
| | Team | 0.003 | 0.004 | 0.347 | 0.729 |
| | Sets × Team | -0.003 | 0.004 | 0.276 | 0.720 |
| FPN-LIMB | Sets | -0.002 | 0.005 | 0.530 | 0.847 |

| | | | | | |
|---|---|---|---|---|---|
| | Team | 0.005 | 0.003 | 0.036 | 0.192 |
| | Sets × Team | < 0.001 | 0.003 | 0.962 | 0.974 |
| FPN-SMN | Sets | -0.002 | 0.004 | 0.566 | 0.865 |
| | Team | -0.001 | 0.003 | 0.525 | 0.847 |
| | Sets × Team | < 0.001 | 0.003 | 0.974 | 0.974 |
| FPN-VAN | Sets | 0.001 | 0.005 | 0.791 | 0.923 |
| | Team | 0.005 | 0.003 | 0.015 | 0.115 |
| | Sets × Team | -0.002 | 0.003 | 0.386 | 0.732 |
| FPN-VIS | Sets | 0.001 | 0.005 | 0.875 | 0.941 |
| | Team | 0.005 | 0.003 | 0.018 | 0.115 |
| | Sets × Team | -0.002 | 0.003 | 0.363 | 0.729 |
| LIMB-LIMB | Sets | -0.006 | 0.010 | 0.364 | 0.729 |
| | Team | -0.003 | 0.006 | 0.564 | 0.865 |
| | Sets × Team | 0.001 | 0.006 | 0.739 | 0.923 |
| LIMB-SMN | Sets | -0.008 | 0.007 | 0.089 | 0.340 |
| | Team | -0.001 | 0.004 | 0.636 | 0.901 |
| | Sets × Team | 0.004 | 0.004 | 0.192 | 0.623 |
| LIMB-VAN | Sets | -0.010 | 0.007 | 0.044 | 0.201 |
| | Team | 0.004 | 0.005 | 0.219 | 0.672 |
| | Sets × Team | 0.006 | 0.004 | 0.073 | 0.294 |
| LIMB-VIS | Sets | 0.002 | 0.004 | 0.514 | 0.847 |
| | Team | -0.001 | 0.003 | 0.752 | 0.923 |
| | Sets × Team | -0.001 | 0.003 | 0.700 | 0.923 |

| | | | | | |
|---|---|---|---|---|---|
| SMN-SMN | Sets | -0.004 | 0.006 | 0.301 | 0.722 |
| | Team | 0.003 | 0.004 | 0.243 | 0.681 |
| | Sets × Team | 0.003 | 0.004 | 0.318 | 0.722 |
| SMN-VAN | Sets | -0.001 | 0.005 | 0.802 | 0.923 |
| | Team | -0.002 | 0.003 | 0.268 | 0.720 |
| | Sets × Team | 0.003 | 0.003 | 0.232 | 0.672 |
| SMN-VIS | Sets | 0.003 | 0.004 | 0.282 | 0.720 |
| | Team | 0.003 | 0.003 | 0.123 | 0.450 |
| | Sets × Team | -0.001 | 0.003 | 0.489 | 0.838 |
| VAN-VAN | Sets | -0.011 | 0.008 | 0.043 | 0.201 |
| | Team | -0.003 | 0.005 | 0.438 | 0.783 |
| | Sets × Team | 0.005 | 0.005 | 0.147 | 0.516 |
| VAN-VIS | Sets | 0.007 | 0.004 | 0.034 | 0.192 |
| | Team | 0.002 | 0.003 | 0.304 | 0.722 |
| | Sets × Team | -0.005 | 0.003 | 0.009 | 0.088 |
| VIS-VIS | Sets | 0.001 | 0.005 | 0.795 | 0.923 |
| | Team | 0.001 | 0.003 | 0.743 | 0.923 |
| | Sets × Team | -0.001 | 0.003 | 0.727 | 0.923 |

**Table S3-2** Full linear mixed-effects model results for associations between network-level functional connectivity differences and Sets, Team, and Sets × Team in the resting-state condition, controlling for Position and Scores.

| Brain network(s) | Predictor | $\beta$ | SE | P | P (FDR-corrected) |
|---|---|---|---|---|---|
| DAN-DAN | Sets | 0.001 | 0.006 | 0.810 | 0.988 |
| | Team | < 0.001 | 0.004 | 0.984 | 0.998 |
| | Sets × Team | -0.006 | 0.005 | 0.088 | 0.490 |
| | Position | < 0.001 | 0.003 | 0.850 | 0.988 |
| | Scores | -0.002 | 0.004 | 0.580 | 0.953 |
| DAN-DMN | Sets | < 0.001 | 0.004 | 0.897 | 0.988 |
| | Team | -0.003 | 0.003 | 0.146 | 0.627 |
| | Sets × Team | -0.002 | 0.003 | 0.289 | 0.811 |
| | Position | 0.001 | 0.002 | 0.520 | 0.950 |
| | Scores | 0.001 | 0.002 | 0.725 | 0.981 |
| DAN-FPN | Sets | 0.001 | 0.006 | 0.846 | 0.988 |
| | Team | -0.001 | 0.004 | 0.827 | 0.988 |
| | Sets × Team | -0.004 | 0.004 | 0.163 | 0.665 |
| | Position | -0.001 | 0.003 | 0.700 | 0.981 |
| | Scores | -0.001 | 0.004 | 0.803 | 0.988 |
| DAN-LIMB | Sets | -0.007 | 0.006 | 0.075 | 0.480 |
| | Team | -0.002 | 0.003 | 0.453 | 0.942 |
| | Sets × Team | 0.001 | 0.004 | 0.675 | 0.981 |
| | Position | -0.001 | 0.003 | 0.504 | 0.942 |
| | Scores | 0.004 | 0.003 | 0.074 | 0.480 |

| | | | | | |
|---|---|---|---|---|---|
| DAN-SMN | Sets | -0.004 | 0.004 | 0.229 | 0.735 |
| | Team | -0.001 | 0.003 | 0.791 | 0.988 |
| | Sets × Team | 0.001 | 0.003 | 0.734 | 0.981 |
| | Position | < 0.001 | 0.002 | 0.877 | 0.988 |
| | Scores | 0.002 | 0.003 | 0.334 | 0.848 |
| DAN-VAN | Sets | 0.002 | 0.005 | 0.545 | 0.953 |
| | Team | -0.002 | 0.003 | 0.319 | 0.848 |
| | Sets × Team | -0.001 | 0.004 | 0.721 | 0.981 |
| | Position | < 0.001 | 0.003 | 0.976 | 0.998 |
| | Scores | -0.002 | 0.003 | 0.393 | 0.888 |
| DAN-VIS | Sets | -0.002 | 0.004 | 0.599 | 0.953 |
| | Team | 0.005 | 0.003 | 0.009 | 0.148 |
| | Sets × Team | -0.002 | 0.003 | 0.280 | 0.811 |
| | Position | < 0.001 | 0.002 | 0.957 | 0.998 |
| | Scores | 0.001 | 0.002 | 0.742 | 0.981 |
| DMN-DMN | Sets | 0.008 | 0.004 | 0.004 | 0.144 |
| | Team | 0.004 | 0.003 | 0.022 | 0.210 |
| | Sets × Team | < 0.001 | 0.003 | 0.972 | 0.998 |
| | Position | < 0.001 | 0.002 | 0.890 | 0.988 |
| | Scores | -0.005 | 0.003 | 0.009 | 0.148 |
| DMN-FPN | Sets | 0.003 | 0.004 | 0.415 | 0.923 |
| | Team | 0.005 | 0.003 | 0.008 | 0.148 |
| | Sets × Team | -0.004 | 0.003 | 0.143 | 0.627 |

| | | | | | |
|---|---|---|---|---|---|
| | Position | 0.001 | 0.002 | 0.529 | 0.950 |
| | Scores | -0.002 | 0.003 | 0.436 | 0.942 |
| DMN-LIMB | Sets | 0.010 | 0.006 | 0.018 | 0.193 |
| | Team | 0.007 | 0.004 | 0.009 | 0.148 |
| | Sets × Team | -0.001 | 0.005 | 0.677 | 0.981 |
| | Position | < 0.001 | 0.003 | 0.853 | 0.988 |
| | Scores | -0.006 | 0.004 | 0.016 | 0.193 |
| DMN-SMN | Sets | 0.001 | 0.004 | 0.717 | 0.981 |
| | Team | < 0.001 | 0.002 | 0.992 | 0.998 |
| | Sets × Team | -0.003 | 0.003 | 0.181 | 0.668 |
| | Position | 0.001 | 0.002 | 0.562 | 0.953 |
| | Scores | -0.001 | 0.002 | 0.468 | 0.942 |
| DMN-VAN | Sets | < 0.001 | 0.004 | 0.883 | 0.988 |
| | Team | < 0.001 | 0.003 | 0.891 | 0.988 |
| | Sets × Team | -0.002 | 0.003 | 0.498 | 0.942 |
| | Position | < 0.001 | 0.003 | 0.954 | 0.998 |
| | Scores | -0.001 | 0.003 | 0.649 | 0.981 |
| DMN-VIS | Sets | 0.007 | 0.004 | 0.014 | 0.193 |
| | Team | 0.007 | 0.002 | < 0.001 | **0.011** |
| | Sets × Team | -0.003 | 0.003 | 0.147 | 0.627 |
| | Position | 0.001 | 0.002 | 0.575 | 0.953 |
| | Scores | -0.005 | 0.002 | 0.003 | 0.139 |
| FPN-FPN | Sets | 0.002 | 0.007 | 0.690 | 0.981 |

| | | | | | |
|---|---|---|---|---|---|
| | Team | 0.003 | 0.004 | 0.360 | 0.855 |
| | Sets × Team | -0.002 | 0.006 | 0.673 | 0.981 |
| | Position | < 0.001 | 0.004 | 0.997 | 0.998 |
| | Scores | -0.003 | 0.004 | 0.280 | 0.811 |
| FPN-LIMB | Sets | -0.003 | 0.005 | 0.481 | 0.942 |
| | Team | 0.005 | 0.003 | 0.034 | 0.279 |
| | Sets × Team | 0.002 | 0.004 | 0.588 | 0.953 |
| | Position | 0.001 | 0.003 | 0.669 | 0.981 |
| | Scores | < 0.001 | 0.003 | 0.990 | 0.998 |
| FPN-SMN | Sets | -0.001 | 0.004 | 0.783 | 0.988 |
| | Team | -0.001 | 0.003 | 0.488 | 0.942 |
| | Sets × Team | -0.001 | 0.003 | 0.645 | 0.981 |
| | Position | -0.002 | 0.002 | 0.245 | 0.748 |
| | Scores | < 0.001 | 0.003 | 0.846 | 0.988 |
| FPN-VAN | Sets | < 0.001 | 0.005 | 0.941 | 0.998 |
| | Team | 0.005 | 0.003 | 0.016 | 0.193 |
| | Sets × Team | -0.003 | 0.004 | 0.369 | 0.862 |
| | Position | 0.001 | 0.003 | 0.473 | 0.942 |
| | Scores | -0.002 | 0.003 | 0.441 | 0.942 |
| FPN-VIS | Sets | 0.001 | 0.005 | 0.764 | 0.988 |
| | Team | 0.005 | 0.003 | 0.020 | 0.206 |
| | Sets × Team | -0.002 | 0.004 | 0.555 | 0.953 |
| | Position | -0.001 | 0.003 | 0.526 | 0.950 |

| | | | | | |
|---|---|---|---|---|---|
| | Scores | -0.002 | 0.003 | 0.323 | 0.848 |
| LIMB-LIMB | Sets | -0.006 | 0.010 | 0.379 | 0.871 |
| | Team | -0.003 | 0.006 | 0.565 | 0.953 |
| | Sets × Team | < 0.001 | 0.008 | 0.998 | 0.998 |
| | Position | < 0.001 | 0.005 | 0.994 | 0.998 |
| | Scores | 0.001 | 0.006 | 0.742 | 0.981 |
| LIMB-SMN | Sets | -0.008 | 0.007 | 0.090 | 0.490 |
| | Team | -0.002 | 0.004 | 0.594 | 0.953 |
| | Sets × Team | -0.005 | 0.005 | 0.220 | 0.735 |
| | Position | < 0.001 | 0.004 | 0.892 | 0.988 |
| | Scores | 0.004 | 0.004 | 0.193 | 0.677 |
| LIMB-VAN | Sets | -0.010 | 0.007 | 0.054 | 0.405 |
| | Team | 0.004 | 0.005 | 0.231 | 0.735 |
| | Sets × Team | -0.002 | 0.006 | 0.678 | 0.981 |
| | Position | < 0.001 | 0.004 | 0.886 | 0.988 |
| | Scores | 0.006 | 0.004 | 0.079 | 0.485 |
| LIMB-VIS | Sets | 0.003 | 0.005 | 0.335 | 0.848 |
| | Team | -0.001 | 0.003 | 0.711 | 0.981 |
| | Sets × Team | -0.001 | 0.004 | 0.768 | 0.988 |
| | Position | -0.003 | 0.003 | 0.170 | 0.665 |
| | Scores | -0.001 | 0.003 | 0.570 | 0.953 |
| SMN-SMN | Sets | -0.006 | 0.006 | 0.170 | 0.665 |
| | Team | 0.004 | 0.004 | 0.142 | 0.627 |

| | | | | | |
|---|---|---|---|---|---|
| | Sets × Team | 0.013 | 0.005 | < 0.001 | **0.011** |
| | Position | 0.004 | 0.003 | 0.103 | 0.536 |
| | Scores | 0.003 | 0.004 | 0.214 | 0.731 |
| SMN-VAN | Sets | 0.001 | 0.005 | 0.851 | 0.988 |
| | Team | -0.003 | 0.003 | 0.236 | 0.735 |
| | Sets × Team | -0.001 | 0.004 | 0.642 | 0.981 |
| | Position | -0.004 | 0.003 | 0.074 | 0.480 |
| | Scores | 0.002 | 0.003 | 0.343 | 0.848 |
| SMN-VIS | Sets | 0.003 | 0.005 | 0.314 | 0.848 |
| | Team | 0.003 | 0.003 | 0.141 | 0.627 |
| | Sets × Team | -0.003 | 0.004 | 0.180 | 0.668 |
| | Position | < 0.001 | 0.003 | 0.924 | 0.998 |
| | Scores | -0.001 | 0.003 | 0.495 | 0.942 |
| VAN-VAN | Sets | -0.009 | 0.008 | 0.091 | 0.490 |
| | Team | -0.003 | 0.005 | 0.449 | 0.942 |
| | Sets × Team | 0.004 | 0.006 | 0.345 | 0.848 |
| | Position | -0.003 | 0.004 | 0.284 | 0.811 |
| | Scores | 0.004 | 0.005 | 0.193 | 0.677 |
| VAN-VIS | Sets | 0.007 | 0.005 | 0.027 | 0.239 |
| | Team | 0.002 | 0.003 | 0.352 | 0.851 |
| | Sets × Team | -0.004 | 0.004 | 0.145 | 0.627 |
| | Position | -0.001 | 0.003 | 0.494 | 0.942 |
| | Scores | -0.005 | 0.003 | 0.007 | 0.148 |

| | | | | | |
|---|---|---|---|---|---|
| VIS-VIS | Sets | 0.001 | 0.005 | 0.796 | 0.988 |
| | Team | < 0.001 | 0.003 | 0.831 | 0.988 |
| | Sets × Team | -0.005 | 0.004 | 0.037 | 0.293 |
| | Position | < 0.001 | 0.003 | 0.910 | 0.995 |
| | Scores | -0.001 | 0.003 | 0.707 | 0.981 |

**Table S3-3** Full linear mixed-effects model results for associations between network-level functional connectivity differences and Sets, Team, and Sets × Team in the game-viewing condition.

| Brain network(s) | Predictor | $\beta$ | SE | P | P (FDR-corrected) |
|---|---|---|---|---|---|
| DAN-DAN | Sets | -0.001 | 0.005 | 0.851 | 0.920 |
| | Team | -0.003 | 0.003 | 0.223 | 0.582 |
| | Sets × Team | 0.002 | 0.003 | 0.243 | 0.582 |
| DAN-DMN | Sets | -0.003 | 0.003 | 0.246 | 0.582 |
| | Team | -0.001 | 0.002 | 0.623 | 0.794 |
| | Sets × Team | 0.003 | 0.002 | 0.054 | 0.582 |
| DAN-FPN | Sets | -0.001 | 0.004 | 0.594 | 0.780 |
| | Team | 0.004 | 0.002 | 0.036 | 0.582 |
| | Sets × Team | 0.001 | 0.002 | 0.452 | 0.695 |
| DAN-LIMB | Sets | -0.001 | 0.005 | 0.766 | 0.858 |
| | Team | -0.003 | 0.003 | 0.168 | 0.582 |
| | Sets × Team | 0.001 | 0.003 | 0.711 | 0.818 |
| DAN-SMN | Sets | < 0.001 | 0.003 | 0.862 | 0.920 |
| | Team | -0.001 | 0.002 | 0.445 | 0.695 |
| | Sets × Team | < 0.001 | 0.002 | 0.986 | 0.987 |
| DAN-VAN | Sets | -0.003 | 0.004 | 0.227 | 0.582 |
| | Team | -0.001 | 0.003 | 0.545 | 0.757 |
| | Sets × Team | 0.003 | 0.003 | 0.112 | 0.582 |
| DAN-VIS | Sets | < 0.001 | 0.005 | 0.897 | 0.931 |
| | Team | -0.003 | 0.003 | 0.169 | 0.582 |

| | | | | | |
|---|---|---|---|---|---|
| | Sets × Team | < 0.001 | 0.003 | 0.872 | 0.920 |
| DMN-DMN | Sets | -0.002 | 0.005 | 0.636 | 0.797 |
| | Team | -0.002 | 0.003 | 0.298 | 0.582 |
| | Sets × Team | 0.001 | 0.003 | 0.660 | 0.804 |
| DMN-FPN | Sets | -0.006 | 0.006 | 0.129 | 0.582 |
| | Team | -0.003 | 0.004 | 0.334 | 0.625 |
| | Sets × Team | 0.003 | 0.004 | 0.287 | 0.582 |
| DMN-LIMB | Sets | -0.009 | 0.006 | 0.040 | 0.582 |
| | Team | 0.001 | 0.004 | 0.686 | 0.812 |
| | Sets × Team | 0.005 | 0.004 | 0.074 | 0.582 |
| DMN-SMN | Sets | -0.006 | 0.004 | 0.024 | 0.582 |
| | Team | -0.001 | 0.003 | 0.681 | 0.812 |
| | Sets × Team | 0.003 | 0.002 | 0.102 | 0.582 |
| DMN-VAN | Sets | -0.006 | 0.006 | 0.107 | 0.582 |
| | Team | -0.001 | 0.004 | 0.557 | 0.757 |
| | Sets × Team | 0.003 | 0.003 | 0.179 | 0.582 |
| DMN-VIS | Sets | -0.004 | 0.004 | 0.176 | 0.582 |
| | Team | -0.002 | 0.003 | 0.344 | 0.628 |
| | Sets × Team | 0.002 | 0.003 | 0.283 | 0.582 |
| FPN-FPN | Sets | -0.005 | 0.008 | 0.365 | 0.647 |
| | Team | -0.004 | 0.005 | 0.265 | 0.582 |
| | Sets × Team | 0.004 | 0.005 | 0.262 | 0.582 |
| FPN-LIMB | Sets | 0.008 | 0.006 | 0.058 | 0.582 |

| | | | | | |
|---|---|---|---|---|---|
| | Team | -0.003 | 0.004 | 0.233 | 0.582 |
| | Sets × Team | -0.005 | 0.004 | 0.065 | 0.582 |
| FPN-SMN | Sets | -0.004 | 0.005 | 0.235 | 0.582 |
| | Team | -0.002 | 0.003 | 0.253 | 0.582 |
| | Sets × Team | 0.002 | 0.003 | 0.369 | 0.647 |
| FPN-VAN | Sets | -0.005 | 0.006 | 0.304 | 0.582 |
| | Team | -0.004 | 0.004 | 0.176 | 0.582 |
| | Sets × Team | 0.002 | 0.004 | 0.403 | 0.658 |
| FPN-VIS | Sets | -0.005 | 0.004 | 0.111 | 0.582 |
| | Team | -0.003 | 0.003 | 0.122 | 0.582 |
| | Sets × Team | 0.002 | 0.003 | 0.272 | 0.582 |
| LIMB-LIMB | Sets | -0.006 | 0.008 | 0.294 | 0.582 |
| | Team | 0.002 | 0.005 | 0.481 | 0.723 |
| | Sets × Team | 0.004 | 0.005 | 0.220 | 0.582 |
| LIMB-SMN | Sets | -0.007 | 0.005 | 0.067 | 0.582 |
| | Team | < 0.001 | 0.003 | 0.987 | 0.987 |
| | Sets × Team | 0.003 | 0.003 | 0.196 | 0.582 |
| LIMB-VAN | Sets | -0.003 | 0.006 | 0.455 | 0.695 |
| | Team | -0.002 | 0.004 | 0.404 | 0.658 |
| | Sets × Team | 0.001 | 0.004 | 0.653 | 0.804 |
| LIMB-VIS | Sets | < 0.001 | 0.004 | 0.876 | 0.920 |
| | Team | -0.001 | 0.003 | 0.528 | 0.757 |
| | Sets × Team | 0.002 | 0.003 | 0.273 | 0.582 |

| | | | | | |
|---|---|---|---|---|---|
| SMN-SMN | Sets | 0.002 | 0.004 | 0.612 | 0.791 |
| | Team | -0.002 | 0.003 | 0.395 | 0.658 |
| | Sets × Team | 0.001 | 0.003 | 0.530 | 0.757 |
| SMN-VAN | Sets | -0.004 | 0.005 | 0.260 | 0.582 |
| | Team | 0.002 | 0.003 | 0.293 | 0.582 |
| | Sets × Team | 0.002 | 0.003 | 0.407 | 0.658 |
| SMN-VIS | Sets | -0.002 | 0.004 | 0.585 | 0.780 |
| | Team | -0.002 | 0.003 | 0.256 | 0.582 |
| | Sets × Team | 0.001 | 0.003 | 0.702 | 0.818 |
| VAN-VAN | Sets | -0.009 | 0.008 | 0.122 | 0.582 |
| | Team | 0.001 | 0.005 | 0.742 | 0.842 |
| | Sets × Team | 0.004 | 0.005 | 0.293 | 0.582 |
| VAN-VIS | Sets | -0.002 | 0.004 | 0.559 | 0.757 |
| | Team | -0.003 | 0.003 | 0.164 | 0.582 |
| | Sets × Team | 0.001 | 0.003 | 0.537 | 0.757 |
| VIS-VIS | Sets | -0.001 | 0.005 | 0.851 | 0.920 |
| | Team | -0.003 | 0.003 | 0.258 | 0.582 |
| | Sets × Team | < 0.001 | 0.003 | 0.984 | 0.987 |

**Table S3-4** Full linear mixed-effects model results for associations between network-level functional connectivity differences and Sets, Team, and Sets × Team in the game-viewing condition, controlling for Position and Scores.

| Brain network(s) | Predictor | *β* | SE | P | P (FDR-corrected) |
|---|---|---|---|---|---|
| DAN-DAN | Sets | 0.001 | 0.005 | 0.816 | 0.894 |
| | Team | -0.003 | 0.003 | 0.185 | 0.663 |
| | Sets × Team | -0.002 | 0.004 | 0.348 | 0.696 |
| | Position | -0.003 | 0.003 | 0.077 | 0.599 |
| | Scores | 0.002 | 0.003 | 0.353 | 0.696 |
| DAN-DMN | Sets | -0.002 | 0.004 | 0.405 | 0.729 |
| | Team | -0.001 | 0.002 | 0.560 | 0.793 |
| | Sets × Team | -0.002 | 0.003 | 0.312 | 0.672 |
| | Position | -0.002 | 0.002 | 0.210 | 0.663 |
| | Scores | 0.003 | 0.002 | 0.081 | 0.599 |
| DAN-FPN | Sets | -0.001 | 0.004 | 0.723 | 0.832 |
| | Team | 0.003 | 0.002 | 0.048 | 0.599 |
| | Sets × Team | -0.004 | 0.003 | 0.072 | 0.599 |
| | Position | -0.001 | 0.002 | 0.456 | 0.742 |
| | Scores | 0.001 | 0.002 | 0.525 | 0.766 |
| DAN-LIMB | Sets | -0.001 | 0.005 | 0.822 | 0.894 |
| | Team | -0.003 | 0.003 | 0.185 | 0.663 |
| | Sets × Team | 0.003 | 0.004 | 0.279 | 0.663 |
| | Position | < 0.001 | 0.003 | 0.823 | 0.894 |
| | Scores | 0.001 | 0.003 | 0.730 | 0.832 |

| | | | | | |
|---|---|---|---|---|---|
| DAN-SMN | Sets | < 0.001 | 0.003 | 0.883 | 0.944 |
| | Team | -0.001 | 0.002 | 0.478 | 0.752 |
| | Sets × Team | 0.002 | 0.003 | 0.316 | 0.672 |
| | Position | < 0.001 | 0.002 | 0.964 | 0.984 |
| | Scores | < 0.001 | 0.002 | 0.986 | 0.986 |
| DAN-VAN | Sets | -0.003 | 0.004 | 0.288 | 0.663 |
| | Team | -0.001 | 0.003 | 0.507 | 0.762 |
| | Sets × Team | -0.002 | 0.003 | 0.398 | 0.729 |
| | Position | -0.001 | 0.002 | 0.619 | 0.805 |
| | Scores | 0.003 | 0.003 | 0.132 | 0.663 |
| DAN-VIS | Sets | -0.001 | 0.005 | 0.721 | 0.832 |
| | Team | -0.003 | 0.003 | 0.199 | 0.663 |
| | Sets × Team | 0.001 | 0.004 | 0.590 | 0.805 |
| | Position | 0.004 | 0.003 | 0.043 | 0.599 |
| | Scores | 0.001 | 0.003 | 0.670 | 0.805 |
| DMN-DMN | Sets | -0.002 | 0.005 | 0.595 | 0.805 |
| | Team | -0.002 | 0.003 | 0.284 | 0.663 |
| | Sets × Team | -0.002 | 0.004 | 0.412 | 0.731 |
| | Position | 0.001 | 0.003 | 0.782 | 0.873 |
| | Scores | 0.001 | 0.003 | 0.642 | 0.805 |
| DMN-FPN | Sets | -0.007 | 0.006 | 0.102 | 0.627 |
| | Team | -0.003 | 0.004 | 0.299 | 0.664 |
| | Sets × Team | -0.006 | 0.005 | 0.070 | 0.599 |

| | | | | | |
|---|---|---|---|---|---|
| | Position | 0.001 | 0.003 | 0.517 | 0.762 |
| | Scores | 0.003 | 0.004 | 0.260 | 0.663 |
| DMN-LIMB | Sets | -0.008 | 0.006 | 0.068 | 0.599 |
| | Team | 0.001 | 0.004 | 0.684 | 0.805 |
| | Sets × Team | 0.002 | 0.005 | 0.607 | 0.805 |
| | Position | -0.002 | 0.003 | 0.442 | 0.736 |
| | Scores | 0.004 | 0.004 | 0.091 | 0.615 |
| DMN-SMN | Sets | -0.008 | 0.004 | 0.008 | 0.301 |
| | Team | -0.001 | 0.003 | 0.637 | 0.805 |
| | Sets × Team | -0.005 | 0.003 | 0.027 | 0.599 |
| | Position | 0.003 | 0.002 | 0.069 | 0.599 |
| | Scores | 0.003 | 0.002 | 0.066 | 0.599 |
| DMN-VAN | Sets | -0.008 | 0.006 | 0.061 | 0.599 |
| | Team | -0.001 | 0.004 | 0.617 | 0.805 |
| | Sets × Team | 0.003 | 0.004 | 0.313 | 0.672 |
| | Position | 0.003 | 0.003 | 0.180 | 0.663 |
| | Scores | 0.004 | 0.004 | 0.133 | 0.663 |
| DMN-VIS | Sets | -0.004 | 0.004 | 0.155 | 0.663 |
| | Team | -0.002 | 0.003 | 0.322 | 0.673 |
| | Sets × Team | -0.003 | 0.003 | 0.250 | 0.663 |
| | Position | 0.001 | 0.002 | 0.666 | 0.805 |
| | Scores | 0.002 | 0.003 | 0.266 | 0.663 |
| FPN-FPN | Sets | -0.008 | 0.008 | 0.126 | 0.663 |

| | | | | | |
|---|---|---|---|---|---|
| | Team | -0.004 | 0.005 | 0.251 | 0.663 |
| | Sets × Team | -0.008 | 0.006 | 0.073 | 0.599 |
| | Position | 0.008 | 0.004 | 0.002 | 0.152 |
| | Scores | 0.005 | 0.005 | 0.147 | 0.663 |
| FPN-LIMB | Sets | 0.007 | 0.006 | 0.107 | 0.627 |
| | Team | -0.003 | 0.004 | 0.298 | 0.664 |
| | Sets × Team | 0.007 | 0.005 | 0.030 | 0.599 |
| | Position | 0.002 | 0.003 | 0.260 | 0.663 |
| | Scores | -0.004 | 0.004 | 0.092 | 0.615 |
| FPN-SMN | Sets | -0.004 | 0.005 | 0.214 | 0.663 |
| | Team | -0.003 | 0.003 | 0.218 | 0.663 |
| | Sets × Team | -0.005 | 0.004 | 0.061 | 0.599 |
| | Position | 0.001 | 0.003 | 0.767 | 0.866 |
| | Scores | 0.002 | 0.003 | 0.359 | 0.698 |
| FPN-VAN | Sets | -0.004 | 0.007 | 0.371 | 0.703 |
| | Team | -0.004 | 0.004 | 0.186 | 0.663 |
| | Sets × Team | 0.003 | 0.005 | 0.427 | 0.732 |
| | Position | -0.001 | 0.003 | 0.680 | 0.805 |
| | Scores | 0.002 | 0.004 | 0.434 | 0.732 |
| FPN-VIS | Sets | -0.005 | 0.005 | 0.107 | 0.627 |
| | Team | -0.003 | 0.003 | 0.119 | 0.663 |
| | Sets × Team | -0.001 | 0.004 | 0.586 | 0.805 |
| | Position | < 0.001 | 0.002 | 0.786 | 0.873 |

| | | | | | |
|---|---|---|---|---|---|
| | Scores | 0.002 | 0.003 | 0.264 | 0.663 |
| LIMB-LIMB | Sets | -0.006 | 0.008 | 0.268 | 0.663 |
| | Team | 0.002 | 0.005 | 0.511 | 0.762 |
| | Sets × Team | -0.005 | 0.006 | 0.282 | 0.663 |
| | Position | 0.001 | 0.004 | 0.731 | 0.832 |
| | Scores | 0.004 | 0.005 | 0.210 | 0.663 |
| LIMB-SMN | Sets | -0.007 | 0.005 | 0.077 | 0.599 |
| | Team | < 0.001 | 0.003 | 0.977 | 0.984 |
| | Sets × Team | -0.002 | 0.004 | 0.423 | 0.732 |
| | Position | < 0.001 | 0.003 | 0.952 | 0.980 |
| | Scores | 0.003 | 0.003 | 0.205 | 0.663 |
| LIMB-VAN | Sets | -0.003 | 0.006 | 0.452 | 0.742 |
| | Team | -0.002 | 0.004 | 0.429 | 0.732 |
| | Sets × Team | 0.002 | 0.005 | 0.497 | 0.762 |
| | Position | < 0.001 | 0.003 | 0.892 | 0.944 |
| | Scores | 0.001 | 0.004 | 0.641 | 0.805 |
| LIMB-VIS | Sets | < 0.001 | 0.005 | 0.897 | 0.944 |
| | Team | -0.001 | 0.003 | 0.556 | 0.793 |
| | Sets × Team | 0.004 | 0.004 | 0.167 | 0.663 |
| | Position | -0.002 | 0.002 | 0.247 | 0.663 |
| | Scores | 0.002 | 0.003 | 0.342 | 0.696 |
| SMN-SMN | Sets | 0.003 | 0.005 | 0.406 | 0.729 |
| | Team | -0.001 | 0.003 | 0.476 | 0.752 |

| | | | | | |
|---|---|---|---|---|---|
| | Sets × Team | 0.008 | 0.004 | 0.001 | 0.122 |
| | Position | -0.002 | 0.003 | 0.222 | 0.663 |
| | Scores | 0.001 | 0.003 | 0.627 | 0.805 |
| SMN-VAN | Sets | -0.004 | 0.005 | 0.235 | 0.663 |
| | Team | 0.002 | 0.003 | 0.278 | 0.663 |
| | Sets × Team | 0.001 | 0.004 | 0.665 | 0.805 |
| | Position | 0.001 | 0.003 | 0.672 | 0.805 |
| | Scores | 0.002 | 0.003 | 0.379 | 0.709 |
| SMN-VIS | Sets | -0.002 | 0.004 | 0.533 | 0.769 |
| | Team | -0.002 | 0.003 | 0.267 | 0.663 |
| | Sets × Team | < 0.001 | 0.003 | 0.859 | 0.925 |
| | Position | 0.001 | 0.002 | 0.684 | 0.805 |
| | Scores | 0.001 | 0.003 | 0.666 | 0.805 |
| VAN-VAN | Sets | -0.007 | 0.008 | 0.246 | 0.663 |
| | Team | 0.002 | 0.005 | 0.663 | 0.805 |
| | Sets × Team | 0.013 | 0.006 | 0.004 | 0.203 |
| | Position | -0.004 | 0.004 | 0.159 | 0.663 |
| | Scores | 0.003 | 0.005 | 0.368 | 0.703 |
| VAN-VIS | Sets | -0.002 | 0.004 | 0.473 | 0.752 |
| | Team | -0.003 | 0.003 | 0.175 | 0.663 |
| | Sets × Team | < 0.001 | 0.003 | 0.912 | 0.953 |
| | Position | 0.001 | 0.002 | 0.511 | 0.762 |
| | Scores | 0.001 | 0.003 | 0.490 | 0.762 |

| | | | | | |
|---|---|---|---|---|---|
| VIS-VIS | Sets | < 0.001 | 0.005 | 0.945 | 0.980 |
| | Team | -0.002 | 0.003 | 0.275 | 0.663 |
| | Sets × Team | 0.003 | 0.004 | 0.344 | 0.696 |
| | Position | -0.001 | 0.003 | 0.669 | 0.805 |
| | Scores | < 0.001 | 0.003 | 0.976 | 0.984 |

**Table S4-1** Full linear mixed-effects model results for associations between network-level functional connectivity differences and Scores, Team, and Scores × Team in the resting-state condition.

| Brain network(s) | Predictor | *β* | SE | P | P (FDR-corrected) |
|---|---|---|---|---|---|
| DAN-DAN | Scores | 0.002 | 0.006 | 0.637 | 0.849 |
| | Team | < 0.001 | 0.004 | 0.962 | 1.000 |
| | Scores × Team | -0.003 | 0.004 | 0.345 | 0.681 |
| DAN-DMN | Scores | < 0.001 | 0.004 | 0.872 | 0.966 |
| | Team | -0.002 | 0.003 | 0.171 | 0.614 |
| | Scores × Team | < 0.001 | 0.002 | 0.791 | 0.927 |
| DAN-FPN | Scores | -0.002 | 0.006 | 0.568 | 0.769 |
| | Team | < 0.001 | 0.004 | 0.886 | 0.966 |
| | Scores × Team | 0.001 | 0.003 | 0.794 | 0.927 |
| DAN-LIMB | Scores | -0.004 | 0.006 | 0.252 | 0.614 |
| | Team | -0.002 | 0.003 | 0.463 | 0.721 |
| | Scores × Team | 0.002 | 0.003 | 0.455 | 0.721 |
| DAN-SMN | Scores | < 0.001 | 0.004 | 0.949 | 1.000 |
| | Team | -0.001 | 0.003 | 0.773 | 0.927 |
| | Scores × Team | < 0.001 | 0.003 | 0.995 | 1.000 |
| DAN-VAN | Scores | -0.002 | 0.005 | 0.558 | 0.768 |
| | Team | -0.002 | 0.003 | 0.334 | 0.681 |
| | Scores × Team | 0.001 | 0.003 | 0.538 | 0.753 |
| DAN-VIS | Scores | 0.004 | 0.004 | 0.191 | 0.614 |
| | Team | 0.005 | 0.002 | 0.010 | 0.197 |

| | | | | | |
|---|---|---|---|---|---|
| | Scores × Team | -0.003 | 0.002 | 0.067 | 0.471 |
| DMN-DMN | Scores | 0.004 | 0.004 | 0.125 | 0.614 |
| | Team | 0.004 | 0.003 | 0.027 | 0.300 |
| | Scores × Team | -0.003 | 0.002 | 0.148 | 0.614 |
| DMN-FPN | Scores | -0.004 | 0.004 | 0.181 | 0.614 |
| | Team | 0.005 | 0.003 | 0.005 | 0.197 |
| | Scores × Team | 0.004 | 0.003 | 0.056 | 0.471 |
| DMN-LIMB | Scores | -0.005 | 0.006 | 0.200 | 0.614 |
| | Team | 0.007 | 0.004 | 0.007 | 0.197 |
| | Scores × Team | 0.004 | 0.004 | 0.109 | 0.611 |
| DMN-SMN | Scores | -0.003 | 0.004 | 0.339 | 0.681 |
| | Team | < 0.001 | 0.002 | 0.908 | 0.978 |
| | Scores × Team | 0.002 | 0.002 | 0.204 | 0.614 |
| DMN-VAN | Scores | -0.003 | 0.004 | 0.282 | 0.644 |
| | Team | < 0.001 | 0.003 | 0.836 | 0.962 |
| | Scores × Team | 0.002 | 0.003 | 0.255 | 0.614 |
| DMN-VIS | Scores | 0.003 | 0.004 | 0.348 | 0.681 |
| | Team | 0.007 | 0.002 | < 0.001 | **0.014** |
| | Scores × Team | -0.001 | 0.002 | 0.444 | 0.721 |
| FPN-FPN | Scores | 0.003 | 0.007 | 0.500 | 0.725 |
| | Team | 0.003 | 0.004 | 0.388 | 0.694 |
| | Scores × Team | -0.003 | 0.004 | 0.320 | 0.681 |
| FPN-LIMB | Scores | < 0.001 | 0.005 | 1.000 | 1.000 |

| | | | | | |
|---|---|---|---|---|---|
| | Team | 0.005 | 0.003 | 0.038 | 0.358 |
| | Scores × Team | < 0.001 | 0.003 | 0.977 | 1.000 |
| FPN-SMN | Scores | -0.003 | 0.004 | 0.238 | 0.614 |
| | Team | -0.001 | 0.003 | 0.501 | 0.725 |
| | Scores × Team | 0.001 | 0.003 | 0.714 | 0.882 |
| FPN-VAN | Scores | -0.006 | 0.005 | 0.080 | 0.521 |
| | Team | 0.006 | 0.003 | 0.011 | 0.197 |
| | Scores × Team | 0.005 | 0.003 | 0.028 | 0.300 |
| FPN-VIS | Scores | -0.002 | 0.005 | 0.482 | 0.725 |
| | Team | 0.005 | 0.003 | 0.020 | 0.284 |
| | Scores × Team | < 0.001 | 0.003 | 0.874 | 0.966 |
| LIMB-LIMB | Scores | 0.006 | 0.010 | 0.363 | 0.681 |
| | Team | -0.003 | 0.006 | 0.509 | 0.725 |
| | Scores × Team | -0.006 | 0.006 | 0.179 | 0.614 |
| LIMB-SMN | Scores | -0.004 | 0.007 | 0.354 | 0.681 |
| | Team | -0.001 | 0.004 | 0.648 | 0.851 |
| | Scores × Team | 0.002 | 0.004 | 0.484 | 0.725 |
| LIMB-VAN | Scores | -0.004 | 0.007 | 0.461 | 0.721 |
| | Team | 0.004 | 0.005 | 0.217 | 0.614 |
| | Scores × Team | 0.001 | 0.004 | 0.663 | 0.857 |
| LIMB-VIS | Scores | 0.003 | 0.005 | 0.417 | 0.721 |
| | Team | -0.001 | 0.003 | 0.676 | 0.861 |
| | Scores × Team | -0.003 | 0.003 | 0.098 | 0.589 |

| | | | | | |
|---|---|---|---|---|---|
| SMN-SMN | Scores | 0.005 | 0.006 | 0.238 | 0.614 |
| | Team | 0.003 | 0.004 | 0.212 | 0.614 |
| | Scores × Team | < 0.001 | 0.004 | 0.885 | 0.966 |
| SMN-VAN | Scores | -0.005 | 0.005 | 0.133 | 0.614 |
| | Team | -0.002 | 0.003 | 0.283 | 0.644 |
| | Scores × Team | 0.002 | 0.003 | 0.364 | 0.681 |
| SMN-VIS | Scores | 0.004 | 0.005 | 0.233 | 0.614 |
| | Team | 0.003 | 0.003 | 0.136 | 0.614 |
| | Scores × Team | -0.002 | 0.003 | 0.203 | 0.614 |
| VAN-VAN | Scores | -0.008 | 0.008 | 0.157 | 0.614 |
| | Team | -0.003 | 0.005 | 0.435 | 0.721 |
| | Scores × Team | 0.002 | 0.005 | 0.460 | 0.721 |
| VAN-VIS | Scores | 0.004 | 0.005 | 0.254 | 0.614 |
| | Team | 0.002 | 0.003 | 0.373 | 0.682 |
| | Scores × Team | -0.004 | 0.003 | 0.066 | 0.471 |
| VIS-VIS | Scores | < 0.001 | 0.005 | 0.966 | 1.000 |
| | Team | 0.001 | 0.003 | 0.771 | 0.927 |
| | Scores × Team | -0.001 | 0.003 | 0.698 | 0.875 |

**Table S4-2** Full linear mixed-effects model results for associations between network-level functional connectivity differences and Scores, Team, and Scores × Team in the resting-state condition, controlling for Sets and Position.

| Brain network(s) | Predictor | *β* | SE | P | P (FDR-corrected) |
|---|---|---|---|---|---|
| DAN-DAN | Scores | 0.003 | 0.006 | 0.449 | 0.815 |
| | Team | < 0.001 | 0.004 | 0.979 | 0.997 |
| | Scores × Team | -0.006 | 0.005 | 0.084 | 0.650 |
| | Position | -0.001 | 0.003 | 0.515 | 0.815 |
| | Sets | -0.003 | 0.004 | 0.331 | 0.718 |
| DAN-DMN | Scores | < 0.001 | 0.004 | 0.934 | 0.991 |
| | Team | -0.003 | 0.003 | 0.151 | 0.650 |
| | Scores × Team | -0.002 | 0.003 | 0.294 | 0.698 |
| | Position | 0.001 | 0.002 | 0.279 | 0.687 |
| | Sets | < 0.001 | 0.002 | 0.777 | 0.891 |
| DAN-FPN | Scores | -0.002 | 0.006 | 0.689 | 0.862 |
| | Team | -0.001 | 0.004 | 0.834 | 0.923 |
| | Scores × Team | -0.004 | 0.004 | 0.162 | 0.650 |
| | Position | < 0.001 | 0.002 | 0.943 | 0.992 |
| | Sets | 0.001 | 0.003 | 0.803 | 0.907 |
| DAN-LIMB | Scores | -0.004 | 0.006 | 0.269 | 0.686 |
| | Team | -0.002 | 0.003 | 0.481 | 0.815 |
| | Scores × Team | 0.001 | 0.004 | 0.644 | 0.856 |
| | Position | -0.001 | 0.002 | 0.720 | 0.862 |
| | Sets | 0.002 | 0.003 | 0.460 | 0.815 |

| | | | | | |
|---|---|---|---|---|---|
| DAN-SMN | Scores | < 0.001 | 0.004 | 0.974 | 0.997 |
| | Team | < 0.001 | 0.003 | 0.796 | 0.906 |
| | Scores × Team | 0.001 | 0.003 | 0.718 | 0.862 |
| | Position | -0.001 | 0.002 | 0.441 | 0.813 |
| | Sets | < 0.001 | 0.003 | 0.983 | 0.997 |
| DAN-VAN | Scores | -0.002 | 0.005 | 0.641 | 0.856 |
| | Team | -0.002 | 0.003 | 0.329 | 0.718 |
| | Scores × Team | -0.001 | 0.004 | 0.711 | 0.862 |
| | Position | -0.001 | 0.002 | 0.695 | 0.862 |
| | Sets | 0.001 | 0.003 | 0.546 | 0.815 |
| DAN-VIS | Scores | 0.004 | 0.004 | 0.134 | 0.650 |
| | Team | 0.004 | 0.002 | 0.011 | 0.254 |
| | Scores × Team | -0.002 | 0.003 | 0.274 | 0.687 |
| | Position | -0.001 | 0.002 | 0.559 | 0.815 |
| | Sets | -0.003 | 0.002 | 0.064 | 0.564 |
| DMN-DMN | Scores | 0.004 | 0.004 | 0.192 | 0.673 |
| | Team | 0.004 | 0.003 | 0.028 | 0.426 |
| | Scores × Team | < 0.001 | 0.003 | 0.971 | 0.997 |
| | Position | 0.001 | 0.002 | 0.249 | 0.678 |
| | Sets | -0.003 | 0.002 | 0.156 | 0.650 |
| DMN-FPN | Scores | -0.004 | 0.005 | 0.229 | 0.673 |
| | Team | 0.005 | 0.003 | 0.006 | 0.254 |
| | Scores × Team | -0.004 | 0.003 | 0.142 | 0.650 |

| | | | | | |
|---|---|---|---|---|---|
| | Position | < 0.001 | 0.002 | 0.758 | 0.879 |
| | Sets | 0.004 | 0.003 | 0.057 | 0.564 |
| DMN-LIMB | Scores | -0.006 | 0.006 | 0.196 | 0.673 |
| | Team | 0.007 | 0.004 | 0.008 | 0.254 |
| | Scores × Team | -0.001 | 0.005 | 0.652 | 0.856 |
| | Position | 0.001 | 0.002 | 0.615 | 0.856 |
| | Sets | 0.004 | 0.004 | 0.107 | 0.650 |
| DMN-SMN | Scores | -0.002 | 0.004 | 0.501 | 0.815 |
| | Team | < 0.001 | 0.002 | 0.952 | 0.995 |
| | Scores × Team | -0.003 | 0.003 | 0.179 | 0.661 |
| | Position | -0.001 | 0.002 | 0.504 | 0.815 |
| | Sets | 0.002 | 0.002 | 0.213 | 0.673 |
| DMN-VAN | Scores | -0.003 | 0.005 | 0.383 | 0.756 |
| | Team | < 0.001 | 0.003 | 0.855 | 0.928 |
| | Scores × Team | -0.002 | 0.003 | 0.498 | 0.815 |
| | Position | -0.001 | 0.002 | 0.549 | 0.815 |
| | Sets | 0.002 | 0.003 | 0.264 | 0.686 |
| DMN-VIS | Scores | 0.003 | 0.004 | 0.235 | 0.673 |
| | Team | 0.007 | 0.003 | < 0.001 | **0.015** |
| | Scores × Team | -0.003 | 0.003 | 0.130 | 0.650 |
| | Position | -0.001 | 0.002 | 0.539 | 0.815 |
| | Sets | -0.001 | 0.002 | 0.429 | 0.813 |
| FPN-FPN | Scores | 0.005 | 0.007 | 0.340 | 0.722 |

| | | | | | |
|---|---|---|---|---|---|
| | Team | 0.003 | 0.004 | 0.389 | 0.756 |
| | Scores × Team | -0.002 | 0.006 | 0.648 | 0.856 |
| | Position | -0.003 | 0.003 | 0.157 | 0.650 |
| | Sets | -0.003 | 0.004 | 0.303 | 0.701 |
| FPN-LIMB | Scores | 0.001 | 0.005 | 0.813 | 0.911 |
| | Team | 0.005 | 0.003 | 0.034 | 0.426 |
| | Scores × Team | 0.002 | 0.004 | 0.587 | 0.831 |
| | Position | -0.003 | 0.002 | 0.098 | 0.650 |
| | Sets | < 0.001 | 0.003 | 0.994 | 0.998 |
| FPN-SMN | Scores | -0.003 | 0.004 | 0.360 | 0.753 |
| | Team | -0.001 | 0.003 | 0.497 | 0.815 |
| | Scores × Team | -0.001 | 0.003 | 0.644 | 0.856 |
| | Position | -0.001 | 0.002 | 0.292 | 0.698 |
| | Sets | 0.001 | 0.003 | 0.735 | 0.872 |
| FPN-VAN | Scores | -0.005 | 0.005 | 0.171 | 0.650 |
| | Team | 0.006 | 0.003 | 0.012 | 0.254 |
| | Scores × Team | -0.003 | 0.004 | 0.369 | 0.754 |
| | Position | -0.002 | 0.002 | 0.151 | 0.650 |
| | Sets | 0.005 | 0.003 | 0.031 | 0.426 |
| FPN-VIS | Scores | -0.001 | 0.005 | 0.708 | 0.862 |
| | Team | 0.005 | 0.003 | 0.021 | 0.367 |
| | Scores × Team | -0.002 | 0.004 | 0.542 | 0.815 |
| | Position | -0.002 | 0.002 | 0.156 | 0.650 |

| | | | | | |
|---|---|---|---|---|---|
| | Sets | < 0.001 | 0.003 | 0.905 | 0.967 |
| LIMB-LIMB | Scores | 0.008 | 0.010 | 0.251 | 0.678 |
| | Team | -0.003 | 0.006 | 0.528 | 0.815 |
| | Scores × Team | < 0.001 | 0.008 | 0.998 | 0.998 |
| | Position | -0.004 | 0.004 | 0.150 | 0.650 |
| | Sets | -0.006 | 0.006 | 0.170 | 0.650 |
| LIMB-SMN | Scores | -0.003 | 0.007 | 0.565 | 0.815 |
| | Team | -0.001 | 0.004 | 0.620 | 0.856 |
| | Scores × Team | -0.005 | 0.005 | 0.234 | 0.673 |
| | Position | -0.002 | 0.003 | 0.225 | 0.673 |
| | Sets | 0.002 | 0.004 | 0.509 | 0.815 |
| LIMB-VAN | Scores | -0.003 | 0.007 | 0.585 | 0.831 |
| | Team | 0.004 | 0.005 | 0.221 | 0.673 |
| | Scores × Team | -0.002 | 0.006 | 0.709 | 0.862 |
| | Position | -0.002 | 0.003 | 0.438 | 0.813 |
| | Sets | 0.001 | 0.004 | 0.679 | 0.862 |
| LIMB-VIS | Scores | 0.002 | 0.005 | 0.517 | 0.815 |
| | Team | -0.001 | 0.003 | 0.654 | 0.856 |
| | Scores × Team | -0.001 | 0.004 | 0.746 | 0.878 |
| | Position | 0.001 | 0.002 | 0.305 | 0.701 |
| | Sets | -0.003 | 0.003 | 0.102 | 0.650 |
| SMN-SMN | Scores | 0.004 | 0.006 | 0.377 | 0.754 |
| | Team | 0.004 | 0.004 | 0.143 | 0.650 |

| | | | | | |
|---|---|---|---|---|---|
| | Scores × Team | 0.013 | 0.005 | < 0.001 | **0.015** |
| | Position | -0.001 | 0.003 | 0.558 | 0.815 |
| | Sets | < 0.001 | 0.004 | 0.899 | 0.967 |
| SMN-VAN | Scores | -0.007 | 0.005 | 0.063 | 0.564 |
| | Team | -0.002 | 0.003 | 0.256 | 0.678 |
| | Scores × Team | -0.001 | 0.004 | 0.663 | 0.860 |
| | Position | 0.004 | 0.002 | 0.012 | 0.254 |
| | Sets | 0.002 | 0.003 | 0.333 | 0.718 |
| SMN-VIS | Scores | 0.004 | 0.005 | 0.246 | 0.678 |
| | Team | 0.003 | 0.003 | 0.158 | 0.650 |
| | Scores × Team | -0.003 | 0.004 | 0.171 | 0.650 |
| | Position | 0.001 | 0.002 | 0.374 | 0.754 |
| | Sets | -0.002 | 0.003 | 0.207 | 0.673 |
| VAN-VAN | Scores | -0.007 | 0.008 | 0.200 | 0.673 |
| | Team | -0.002 | 0.005 | 0.475 | 0.815 |
| | Scores × Team | 0.004 | 0.006 | 0.328 | 0.718 |
| | Position | -0.003 | 0.003 | 0.231 | 0.673 |
| | Sets | 0.002 | 0.005 | 0.473 | 0.815 |
| VAN-VIS | Scores | 0.005 | 0.005 | 0.168 | 0.650 |
| | Team | 0.002 | 0.003 | 0.409 | 0.786 |
| | Scores × Team | -0.004 | 0.004 | 0.126 | 0.650 |
| | Position | -0.001 | 0.002 | 0.562 | 0.815 |
| | Sets | -0.004 | 0.003 | 0.063 | 0.564 |

| | | | | | |
|---|---|---|---|---|---|
| VIS-VIS | Scores | 0.001 | 0.005 | 0.760 | 0.879 |
| | Team | < 0.001 | 0.003 | 0.847 | 0.927 |
| | Scores × Team | -0.005 | 0.004 | 0.036 | 0.426 |
| | Position | < 0.001 | 0.002 | 0.837 | 0.923 |
| | Sets | -0.001 | 0.003 | 0.683 | 0.862 |

**Table S4-3** Full linear mixed-effects model results for associations between network-level functional connectivity differences and Scores, Team, and Scores × Team in the game-viewing condition.

| Brain network(s) | Predictor | $\beta$ | SE | P | P (FDR-corrected) |
|---|---|---|---|---|---|
| DAN-DAN | Scores | -0.002 | 0.005 | 0.473 | 0.829 |
| | Team | -0.003 | 0.003 | 0.221 | 0.829 |
| | Scores × Team | < 0.001 | 0.003 | 0.996 | 0.997 |
| DAN-DMN | Scores | -0.004 | 0.004 | 0.088 | 0.829 |
| | Team | -0.001 | 0.002 | 0.661 | 0.896 |
| | Scores × Team | 0.002 | 0.002 | 0.212 | 0.829 |
| DAN-FPN | Scores | -0.005 | 0.004 | 0.077 | 0.829 |
| | Team | 0.004 | 0.002 | 0.032 | 0.829 |
| | Scores × Team | 0.002 | 0.002 | 0.161 | 0.829 |
| DAN-LIMB | Scores | 0.002 | 0.005 | 0.555 | 0.829 |
| | Team | -0.003 | 0.003 | 0.161 | 0.829 |
| | Scores × Team | -0.001 | 0.003 | 0.489 | 0.829 |
| DAN-SMN | Scores | 0.001 | 0.003 | 0.546 | 0.829 |
| | Team | -0.001 | 0.002 | 0.426 | 0.829 |
| | Scores × Team | -0.001 | 0.002 | 0.494 | 0.829 |
| DAN-VAN | Scores | -0.002 | 0.004 | 0.487 | 0.829 |
| | Team | -0.001 | 0.003 | 0.560 | 0.829 |
| | Scores × Team | 0.001 | 0.003 | 0.681 | 0.896 |
| DAN-VIS | Scores | 0.003 | 0.005 | 0.416 | 0.829 |
| | Team | -0.003 | 0.003 | 0.196 | 0.829 |

| | | | | | |
|---|---|---|---|---|---|
| | Scores × Team | 0.001 | 0.003 | 0.706 | 0.898 |
| DMN-DMN | Scores | -0.001 | 0.005 | 0.791 | 0.923 |
| | Team | -0.002 | 0.003 | 0.306 | 0.829 |
| | Scores × Team | 0.001 | 0.003 | 0.758 | 0.916 |
| DMN-FPN | Scores | -0.003 | 0.006 | 0.512 | 0.829 |
| | Team | -0.003 | 0.004 | 0.342 | 0.829 |
| | Scores × Team | 0.002 | 0.004 | 0.562 | 0.829 |
| DMN-LIMB | Scores | -0.006 | 0.006 | 0.186 | 0.829 |
| | Team | 0.001 | 0.004 | 0.683 | 0.896 |
| | Scores × Team | 0.002 | 0.004 | 0.376 | 0.829 |
| DMN-SMN | Scores | 0.001 | 0.004 | 0.820 | 0.931 |
| | Team | -0.001 | 0.003 | 0.694 | 0.897 |
| | Scores × Team | < 0.001 | 0.002 | 0.970 | 0.997 |
| DMN-VAN | Scores | -0.003 | 0.006 | 0.419 | 0.829 |
| | Team | -0.001 | 0.004 | 0.616 | 0.877 |
| | Scores × Team | 0.004 | 0.003 | 0.138 | 0.829 |
| DMN-VIS | Scores | -0.001 | 0.004 | 0.680 | 0.896 |
| | Team | -0.002 | 0.003 | 0.350 | 0.829 |
| | Scores × Team | 0.001 | 0.003 | 0.725 | 0.902 |
| FPN-FPN | Scores | 0.002 | 0.008 | 0.649 | 0.896 |
| | Team | -0.004 | 0.005 | 0.300 | 0.829 |
| | Scores × Team | 0.002 | 0.005 | 0.466 | 0.829 |
| FPN-LIMB | Scores | < 0.001 | 0.006 | 0.992 | 0.997 |

| | | | | | |
|---|---|---|---|---|---|
| | Team | -0.003 | 0.004 | 0.270 | 0.829 |
| | Scores × Team | 0.003 | 0.004 | 0.298 | 0.829 |
| FPN-SMN | Scores | -0.001 | 0.005 | 0.730 | 0.902 |
| | Team | -0.002 | 0.003 | 0.252 | 0.829 |
| | Scores × Team | < 0.001 | 0.003 | 0.909 | 0.979 |
| FPN-VAN | Scores | -0.005 | 0.006 | 0.250 | 0.829 |
| | Team | -0.004 | 0.004 | 0.185 | 0.829 |
| | Scores × Team | 0.003 | 0.004 | 0.306 | 0.829 |
| FPN-VIS | Scores | 0.001 | 0.005 | 0.766 | 0.916 |
| | Team | -0.003 | 0.003 | 0.114 | 0.829 |
| | Scores × Team | -0.001 | 0.003 | 0.549 | 0.829 |
| LIMB-LIMB | Scores | -0.006 | 0.008 | 0.245 | 0.829 |
| | Team | 0.003 | 0.005 | 0.439 | 0.829 |
| | Scores × Team | 0.005 | 0.005 | 0.175 | 0.829 |
| LIMB-SMN | Scores | -0.004 | 0.005 | 0.304 | 0.829 |
| | Team | < 0.001 | 0.003 | 0.980 | 0.997 |
| | Scores × Team | 0.002 | 0.003 | 0.489 | 0.829 |
| LIMB-VAN | Scores | -0.005 | 0.006 | 0.304 | 0.829 |
| | Team | -0.002 | 0.004 | 0.429 | 0.829 |
| | Scores × Team | 0.003 | 0.004 | 0.245 | 0.829 |
| LIMB-VIS | Scores | -0.002 | 0.005 | 0.487 | 0.829 |
| | Team | -0.001 | 0.003 | 0.562 | 0.829 |
| | Scores × Team | 0.001 | 0.003 | 0.550 | 0.829 |

| | | | | | |
|---|---|---|---|---|---|
| SMN-SMN | Scores | 0.004 | 0.005 | 0.176 | 0.829 |
| | Team | -0.002 | 0.003 | 0.378 | 0.829 |
| | Scores × Team | -0.003 | 0.003 | 0.139 | 0.829 |
| SMN-VAN | Scores | -0.002 | 0.005 | 0.560 | 0.829 |
| | Team | 0.002 | 0.003 | 0.276 | 0.829 |
| | Scores × Team | 0.002 | 0.003 | 0.445 | 0.829 |
| SMN-VIS | Scores | 0.001 | 0.004 | 0.818 | 0.931 |
| | Team | -0.002 | 0.003 | 0.257 | 0.829 |
| | Scores × Team | < 0.001 | 0.003 | 0.903 | 0.979 |
| VAN-VAN | Scores | -0.004 | 0.008 | 0.517 | 0.829 |
| | Team | 0.001 | 0.005 | 0.774 | 0.916 |
| | Scores × Team | < 0.001 | 0.005 | 0.997 | 0.997 |
| VAN-VIS | Scores | -0.001 | 0.004 | 0.859 | 0.962 |
| | Team | -0.003 | 0.003 | 0.179 | 0.829 |
| | Scores × Team | 0.001 | 0.003 | 0.593 | 0.860 |
| VIS-VIS | Scores | < 0.001 | 0.005 | 0.903 | 0.979 |
| | Team | -0.003 | 0.003 | 0.253 | 0.829 |
| | Scores × Team | < 0.001 | 0.003 | 0.950 | 0.997 |

**Table S4-4** Full linear mixed-effects model results for associations between network-level functional connectivity differences and Scores, Team, and Scores × Team in the game-viewing condition, controlling for Sets and Position.

| Brain network(s) | Predictor | $\beta$ | SE | P | P (FDR-corrected) |
|---|---|---|---|---|---|
| DAN-DAN | Scores | -0.004 | 0.005 | 0.292 | 0.810 |
| | Team | -0.003 | 0.003 | 0.188 | 0.810 |
| | Scores × Team | -0.002 | 0.004 | 0.359 | 0.810 |
| | Position | 0.004 | 0.002 | 0.010 | 0.365 |
| | Sets | < 0.001 | 0.003 | 0.961 | 0.989 |
| DAN-DMN | Scores | -0.005 | 0.004 | 0.059 | 0.755 |
| | Team | -0.001 | 0.002 | 0.608 | 0.843 |
| | Scores × Team | -0.002 | 0.003 | 0.338 | 0.810 |
| | Position | 0.002 | 0.001 | 0.068 | 0.755 |
| | Sets | 0.002 | 0.002 | 0.198 | 0.810 |
| DAN-FPN | Scores | -0.005 | 0.004 | 0.102 | 0.810 |
| | Team | 0.004 | 0.002 | 0.041 | 0.640 |
| | Scores × Team | -0.004 | 0.003 | 0.077 | 0.755 |
| | Position | 0.001 | 0.002 | 0.571 | 0.839 |
| | Sets | 0.002 | 0.002 | 0.160 | 0.810 |
| DAN-LIMB | Scores | 0.001 | 0.005 | 0.667 | 0.849 |
| | Team | -0.003 | 0.003 | 0.177 | 0.810 |
| | Scores × Team | 0.003 | 0.004 | 0.278 | 0.810 |
| | Position | < 0.001 | 0.002 | 0.845 | 0.953 |
| | Sets | -0.001 | 0.003 | 0.497 | 0.810 |

| | | | | | |
|---|---|---|---|---|---|
| DAN-SMN | Scores | 0.001 | 0.004 | 0.582 | 0.839 |
| | Team | -0.001 | 0.002 | 0.459 | 0.810 |
| | Scores × Team | 0.002 | 0.003 | 0.319 | 0.810 |
| | Position | < 0.001 | 0.001 | 0.690 | 0.855 |
| | Sets | -0.001 | 0.002 | 0.492 | 0.810 |
| DAN-VAN | Scores | -0.002 | 0.004 | 0.470 | 0.810 |
| | Team | -0.001 | 0.003 | 0.526 | 0.819 |
| | Scores × Team | -0.002 | 0.003 | 0.419 | 0.810 |
| | Position | 0.001 | 0.002 | 0.454 | 0.810 |
| | Sets | 0.001 | 0.003 | 0.673 | 0.849 |
| DAN-VIS | Scores | 0.003 | 0.005 | 0.470 | 0.810 |
| | Team | -0.003 | 0.003 | 0.206 | 0.810 |
| | Scores × Team | 0.002 | 0.004 | 0.581 | 0.839 |
| | Position | < 0.001 | 0.002 | 0.932 | 0.979 |
| | Sets | 0.001 | 0.003 | 0.702 | 0.862 |
| DMN-DMN | Scores | < 0.001 | 0.005 | 0.891 | 0.965 |
| | Team | -0.002 | 0.003 | 0.291 | 0.810 |
| | Scores × Team | -0.002 | 0.004 | 0.419 | 0.810 |
| | Position | < 0.001 | 0.002 | 0.800 | 0.949 |
| | Sets | 0.001 | 0.003 | 0.768 | 0.920 |
| DMN-FPN | Scores | -0.001 | 0.006 | 0.837 | 0.953 |
| | Team | -0.003 | 0.004 | 0.312 | 0.810 |
| | Scores × Team | -0.006 | 0.005 | 0.074 | 0.755 |

| | | | | | |
|---|---|---|---|---|---|
| | Position | -0.003 | 0.002 | 0.138 | 0.810 |
| | Sets | 0.001 | 0.004 | 0.596 | 0.839 |
| DMN-LIMB | Scores | -0.005 | 0.006 | 0.215 | 0.810 |
| | Team | 0.001 | 0.004 | 0.654 | 0.849 |
| | Scores × Team | 0.002 | 0.005 | 0.580 | 0.839 |
| | Position | -0.001 | 0.002 | 0.499 | 0.810 |
| | Sets | 0.002 | 0.004 | 0.383 | 0.810 |
| DMN-SMN | Scores | 0.003 | 0.004 | 0.391 | 0.810 |
| | Team | -0.001 | 0.003 | 0.646 | 0.849 |
| | Scores × Team | -0.005 | 0.003 | 0.030 | 0.584 |
| | Position | -0.003 | 0.002 | 0.020 | 0.464 |
| | Sets | < 0.001 | 0.002 | 0.973 | 0.990 |
| DMN-VAN | Scores | -0.003 | 0.006 | 0.485 | 0.810 |
| | Team | -0.001 | 0.004 | 0.668 | 0.849 |
| | Scores × Team | 0.003 | 0.004 | 0.288 | 0.810 |
| | Position | -0.002 | 0.002 | 0.236 | 0.810 |
| | Sets | 0.004 | 0.003 | 0.143 | 0.810 |
| DMN-VIS | Scores | < 0.001 | 0.004 | 0.903 | 0.965 |
| | Team | -0.002 | 0.003 | 0.332 | 0.810 |
| | Scores × Team | -0.003 | 0.003 | 0.262 | 0.810 |
| | Position | -0.001 | 0.002 | 0.324 | 0.810 |
| | Sets | 0.001 | 0.003 | 0.749 | 0.912 |
| FPN-FPN | Scores | 0.004 | 0.008 | 0.457 | 0.810 |

| | | | | | |
|---|---|---|---|---|---|
| | Team | -0.004 | 0.005 | 0.266 | 0.810 |
| | Scores × Team | -0.008 | 0.006 | 0.080 | 0.755 |
| | Position | -0.001 | 0.003 | 0.639 | 0.849 |
| | Sets | 0.002 | 0.005 | 0.482 | 0.810 |
| FPN-LIMB | Scores | -0.001 | 0.006 | 0.806 | 0.949 |
| | Team | -0.003 | 0.004 | 0.314 | 0.810 |
| | Scores × Team | 0.007 | 0.005 | 0.033 | 0.584 |
| | Position | < 0.001 | 0.002 | 0.841 | 0.953 |
| | Sets | 0.003 | 0.004 | 0.287 | 0.810 |
| FPN-SMN | Scores | < 0.001 | 0.005 | 0.976 | 0.990 |
| | Team | -0.003 | 0.003 | 0.222 | 0.810 |
| | Scores × Team | -0.005 | 0.004 | 0.064 | 0.755 |
| | Position | -0.001 | 0.002 | 0.330 | 0.810 |
| | Sets | < 0.001 | 0.003 | 0.937 | 0.979 |
| FPN-VAN | Scores | -0.005 | 0.007 | 0.255 | 0.810 |
| | Team | -0.004 | 0.004 | 0.202 | 0.810 |
| | Scores × Team | 0.003 | 0.005 | 0.411 | 0.810 |
| | Position | -0.001 | 0.003 | 0.681 | 0.851 |
| | Sets | 0.003 | 0.004 | 0.308 | 0.810 |
| FPN-VIS | Scores | 0.002 | 0.005 | 0.545 | 0.839 |
| | Team | -0.003 | 0.003 | 0.115 | 0.810 |
| | Scores × Team | -0.001 | 0.004 | 0.599 | 0.839 |
| | Position | -0.002 | 0.002 | 0.148 | 0.810 |

| | | | | | |
|---|---|---|---|---|---|
| | Sets | -0.001 | 0.003 | 0.526 | 0.819 |
| LIMB-LIMB | Scores | -0.006 | 0.008 | 0.294 | 0.810 |
| | Team | 0.002 | 0.005 | 0.471 | 0.810 |
| | Scores × Team | -0.005 | 0.006 | 0.299 | 0.810 |
| | Position | < 0.001 | 0.003 | 0.914 | 0.969 |
| | Sets | 0.005 | 0.005 | 0.178 | 0.810 |
| LIMB-SMN | Scores | -0.003 | 0.005 | 0.502 | 0.810 |
| | Team | < 0.001 | 0.003 | 0.993 | 0.997 |
| | Scores × Team | -0.002 | 0.004 | 0.443 | 0.810 |
| | Position | -0.002 | 0.002 | 0.144 | 0.810 |
| | Sets | 0.001 | 0.003 | 0.515 | 0.819 |
| LIMB-VAN | Scores | -0.004 | 0.007 | 0.343 | 0.810 |
| | Team | -0.002 | 0.004 | 0.458 | 0.810 |
| | Scores × Team | 0.003 | 0.005 | 0.483 | 0.810 |
| | Position | -0.001 | 0.003 | 0.441 | 0.810 |
| | Sets | 0.003 | 0.004 | 0.251 | 0.810 |
| LIMB-VIS | Scores | -0.004 | 0.005 | 0.212 | 0.810 |
| | Team | -0.001 | 0.003 | 0.580 | 0.839 |
| | Scores × Team | 0.004 | 0.004 | 0.159 | 0.810 |
| | Position | 0.003 | 0.002 | 0.016 | 0.453 |
| | Sets | 0.001 | 0.003 | 0.503 | 0.810 |
| SMN-SMN | Scores | 0.002 | 0.005 | 0.642 | 0.849 |
| | Team | -0.002 | 0.003 | 0.441 | 0.810 |

| | | | | | |
|---|---|---|---|---|---|
| | Scores × Team | 0.008 | 0.004 | 0.001 | 0.120 |
| | Position | 0.004 | 0.002 | 0.003 | 0.193 |
| | Sets | -0.003 | 0.003 | 0.159 | 0.810 |
| SMN-VAN | Scores | -0.002 | 0.005 | 0.645 | 0.849 |
| | Team | 0.002 | 0.003 | 0.262 | 0.810 |
| | Scores × Team | 0.001 | 0.004 | 0.646 | 0.849 |
| | Position | -0.001 | 0.002 | 0.362 | 0.810 |
| | Sets | 0.002 | 0.003 | 0.456 | 0.810 |
| SMN-VIS | Scores | 0.001 | 0.004 | 0.764 | 0.920 |
| | Team | -0.002 | 0.003 | 0.266 | 0.810 |
| | Scores × Team | < 0.001 | 0.003 | 0.853 | 0.953 |
| | Position | -0.001 | 0.002 | 0.575 | 0.839 |
| | Sets | < 0.001 | 0.003 | 0.893 | 0.965 |
| VAN-VAN | Scores | -0.005 | 0.008 | 0.427 | 0.810 |
| | Team | 0.002 | 0.005 | 0.659 | 0.849 |
| | Scores × Team | 0.013 | 0.006 | 0.004 | 0.193 |
| | Position | -0.002 | 0.003 | 0.408 | 0.810 |
| | Sets | < 0.001 | 0.005 | 0.997 | 0.997 |
| VAN-VIS | Scores | < 0.001 | 0.004 | 0.880 | 0.965 |
| | Team | -0.003 | 0.003 | 0.184 | 0.810 |
| | Scores × Team | < 0.001 | 0.003 | 0.897 | 0.965 |
| | Position | < 0.001 | 0.002 | 0.839 | 0.953 |
| | Sets | 0.001 | 0.003 | 0.597 | 0.839 |

| | | | | | |
|---|---|---|---|---|---|
| VIS-VIS | Scores | -0.001 | 0.005 | 0.857 | 0.953 |
| | Team | -0.002 | 0.003 | 0.275 | 0.810 |
| | Scores × Team | 0.003 | 0.004 | 0.344 | 0.810 |
| | Position | < 0.001 | 0.002 | 0.817 | 0.953 |
| | Sets | < 0.001 | 0.003 | 0.951 | 0.987 |
